# Histone demethylase enzymes KDM5A and KDM5B modulate immune response by suppressing transcription of endogenous retroviral elements

**DOI:** 10.1101/2024.09.23.614494

**Authors:** Huadong Chen, Letitia Sarah, Daniela Pucciarelli, Ying Mao, Morgan E. Diolaiti, Danica Galonić Fujimori, Alan Ashworth

**Affiliations:** Helen Diller Family Comprehensive Cancer Center, University of California, San Francisco, California; Chemistry and Chemical Biology Graduate Program, University of California, San Francisco, California; Department of Cellular and Molecular Pharmacology, University of California, San Francisco, California; Department of Pharmaceutical Chemistry, University of California, San Francisco, California

## Abstract

Epigenetic factors, including lysine-specific demethylases such as the KDM5 paralogs KDM5A and KDM5B have been implicated in cancer and the regulation of immune responses. Here, we performed a comprehensive multiomic study in cells lacking KDM5A or KDM5B to map changes in transcriptional regulation and chromatin organization. RNA-seq analysis revealed a significant decrease in the expression of Krüppel-associated box containing zinc finger (*KRAB-ZNF*) genes in KDM5A or KDM5B knockout cell lines, which was accompanied by changes ATAC-seq and H3K4me3 ChIP-seq. Pharmacological inhibition of KDM5A and KDM5B catalytic activity with a pan-KDM5 inhibitor, CPI-455, did not significantly change KRAB-ZNF expression, raising the possibility that regulation of KRAB-ZNF expression does not require KDM5A and KDM5B demethylase activity. KRAB-ZNF are recognized suppressors of the transcription of endogenous retroviruses (ERVs) and HAP1 cells with *KDM5A* or *KDM5B* gene inactivation showed elevated ERV expression, increased dsRNA levels and elevated levels of immune response genes. Acute degradation of KDM5A using a dTAG system in HAP1 cells led to increased ERV expression, demonstrating that de-repression of ERV genes occurs rapidly after loss of KDM5A. Co-immunoprecipitation of KDM5A revealed an interaction with the Nucleosome Remodeling and Deacetylase (NuRD) complex suggesting that KDM5A and NuRD may act together to regulate the expression of ERVs through KRAB-ZNFs. These findings reveal roles of KDM5A and KDM5B in modulating ERV expression and underscore the therapeutic potential of using degraders of KDM5A and KDM5B to modulate tumor immune responses.

**Author Summary:** The histone demethylases KDM5A and KDM5B are transcriptional repressors that play an important role in cancer and immune response, making them attractive drug targets. Unfortunately, small molecule inhibitors, including CPI-455, that block KDM5A and KDM5B enzymatic activity, have shown only limited effectiveness at suppressing cancer cell viability as single agents in vitro. In this study we undertook a multi-omics approach to map transcriptional and chromatin changes in KDM5A and KDM5B deficient cells compared to those treated with CPI-455. The datasets revealed that KDM5A and KDM5B modulate the expression of KRAB-ZNF genes and that loss of either gene was associated with increased expression of ERV genes and upregulation of immune response markers. Surprisingly, pharmacological inhibition of these enzymes did not phenocopy genetic ablation. In contrast, acute degradation of KDM5A using a dTAG system caused an increase in ERV expression, providing evidence that this immune modulation is independent of demethylase activity. Together with the limited success of small molecule inhibitors, our data provide strong rationale for the development of KDM5A and KDM5B degraders to modulate tumor immune responses.

## Introduction

Epigenetic alterations are common in tumorigenesis, influencing tumor initiation, progression, chemoresistance, and immune regulation and dysregulation of chromatin modifying enzymes can lead to activation of oncogenes or repression of tumor suppressor genes, disrupting critical signaling pathways [1]. Epigenetic regulators also have an important role in immune cell function and antitumor immunity [2]. Consequently, therapies targeting chromatin modifying enzymes, either alone or in combination with immunotherapies, have emerged as a promising strategy to treat a variety of tumors [3–5].

KDM5, a family of histone H3 lysine 4 demethylases, is of interest as a potential therapeutic target. Among the four paralogs of KDM5 (A-D), the genes encoding KDM5A and KDM5B are frequently amplified and overexpressed in several cancers including those of the breast, prostate, liver, lung, stomach, head and neck and those of the nervous system [6]. Additionally, KDM5B contributes to therapeutic resistance in estrogen receptor (ER) positive breast cancer by enhancing transcriptomic heterogeneity [7]. Recent evidence also highlights the role of KDM5 demethylases in immune regulation (7,8). KDM5B and KDM5C have been reported to suppress expression of the stimulator of interferon genes (STING) via the removal of H3K4me3, an active transcription mark antagonized by KDM5 enzymes, at gene promoters [8]; in various human tumor types, KDM5B expression inversely correlates to the expression of STING. In epithelial ovarian cancer, KDM5A regulates CD8^+^ T-cell infiltration by silencing genes associated with the antigen processing and presentation pathway [9]. This regulatory mechanism is counteracted by KDM5A inhibition, suggesting a demethylation-dependent function. Apart from its immune-regulatory demethylase activity, KDM5 proteins also display demethylase-independent functions. In a melanoma mouse model, KDM5B promotes immune evasion through silencing of transposable elements [10]. This is achieved through a KDM5B-mediated scaffolding of a repressive methyltransferase, SETDB1.

Additionally, KDM5B plays a demethylase-independent role in suppressing acute myeloid leukemia (AML) by recruiting HDAC1-containing transcriptional repressive machinery [11]. This results in the downregulation of stemness genes and the suppression of AML growth. Motivated by this emerging understanding of KDM5 demethylation and scaffolding functions in cancer and immunity, we sought to further elucidate the role of KDM5 proteins as chromatin regulators utilizing a multiomics approach. We report that genetic inactivation of *KDM5A* or *KDM5B* in HAP1 cells leads to downregulation of the expression of select Krüppel-associated box containing zinc finger (KRAB-ZNF) genes in a catalysis-independent manner. This downregulation of KRAB-ZNF results in enhanced transcription of endogenous retroviruses (ERVs). Additionally, we show that KDM5A associates with components of Nucleosome Remodeling and Deacetylase (NuRD) complex and KRAB-ZNF repressor complex, implicating KDM5A in the assembly of these complexes. Taken together, our results reveal that KDM5A and KDM5B regulate immune responses by inhibiting ERV expression, nominating KDM5A and KDM5B as potential therapeutic targets for enhancing antitumor immune response.

## Results

### KDM5A/B knockout results in closed chromatin and reduced KRAB-ZNFs expression

To explore the role of KDM5A and KDM5B in regulating chromatin state, we used HAP1 cells in which the *KDM5A* and *KDM5B* genes had been inactivated using CRISPR/Cas9 (HAP1^Δ5A^ and HAP1^Δ5Β^). We first verified disruption of the genes by PCR and DNA sequencing and confirmed loss of KDM5A and KDM5B protein expression by western blotting (Fig. 1A and Fig. S1A). To identify genome-wide transcriptional changes that occur upon loss of *KDM5A* or *KDM5B*, we performed RNA-seq on RNA prepared from HAP1 knockout and parental cells (Fig. 1B and 1C). Compared to HAP1 parental cells, HAP1^Δ5A^ exhibit 811 up-regulated genes (DEGs; log2FC ≥ −1, Padj ≤ 0.05) and 1051 down-regulated genes (DEGs; log2FC ≤ −1, Padj ≤ 0.05), while HAP1^Δ5Β^ cells had 1240 up-regulated genes and 1031 down-regulated genes (Fig. 1B-E). Amongst these up-regulated DEGs, 570 up-regulated genes and 717 down-regulated genes were common to both HAP1^Δ5A^ and HAP1^Δ5Β^ demonstrating considerable functional similarity between KDM5A and KDM5B (Fig. 1D and 1E). Amongst the 717 DEGs shared by the two knockout lines, the pathway (DAVID Bioinformatics Resources 6.8) with the greatest numerical difference was the KRAB zinc-finger protein (KRAB-ZNFs) group of genes. In total, 69 KRAB-ZNFs genes were down-regulated in both HAP1^Δ5A^ and HAP1^Δ5Β^ (Fig. 1E) and amongst the top 16 down-regulated DEGs, 9 were KRAB-ZNFs genes (Table 1), suggesting the importance of KDM5A and KDM5B for the regulation of KRAB-ZNFs. RT-qPCR verified the downregulation of KRAB-ZNFs in HAP1^Δ5A^ and HAP1^Δ5Β^ cells (Fig. 1F). Gene Ontology (GO) pathway analysis of up-regulated DEGs in HAP1^Δ5A^ cells showed enrichment of chemorepellent activity-related genes (*SEMA5A, SEMA4A, EPHA7, SEMA6A, SEMA3C, SEMA3D, SEMA3A, SEMA6D, SEMA3G, NRG1, ENA5, FLRT2, NRG3*) and Calcium Ion-regulated Exocytosis of Neurotransmitter pathway genes (*SYT3, RPH3AL, SYT1, SYT15, C2CD4C, LOC102724488888C, LOC10272448888C 8, DOC2B, SYT7, SYT6, SYTL2, Rims2, Rims1, Rims4, SYT11*). GO pathway analysis of up-regulated DEGs in HAP1^Δ5Β^ cells showed enrichment in genes related to transmembrane receptor protein tyrosine kinase activity (*RET, PDGFRA, NTRK2, FLT1, FLT4, MERTK, ERBB3, AXL, ERBB4, ERBB2, KDR, ROR1, TEK, ROR2, MET, FGFR2*) and Ras guanyl-nucleotide exchange factor activity related genes (*RET, KLB, SHC2, CAMK2D, RASGRF2, PDGFB, ADRB1, RASGRP1, FGF4, FGF5, FGF8, RASGEF1A, ERBB3, ERBB4, RASGEF1B, ERBB2, NEFL, PDGFRB, PDGFRA, ANGPT1, ACTN2, NRG2, SPTB, GRIN2D, GDNF, FGF19, TEK, FGFR4, FGFR3, FGFR2*). KEGG pathway analysis of DEGs revealed upregulation of genes involved in Axon guidance, Neuroactive ligand-receptor interaction, and Cytokine-cytokine receptor interaction pathways in HAP1^Δ5A^. In HAP1^Δ5Β^, mRNAs for genes involved in PI3K-Akt signaling, Axon guidance, and Focal adhesion pathways were elevated (Fig. 1G and H). Comparing the GO-analysis for the top 20 of up-regulated DEGs genes in HAP1^Δ5A^ vs HAP1^Δ5Β^, revealed no overlap (Table 2).

**Fig.1.**
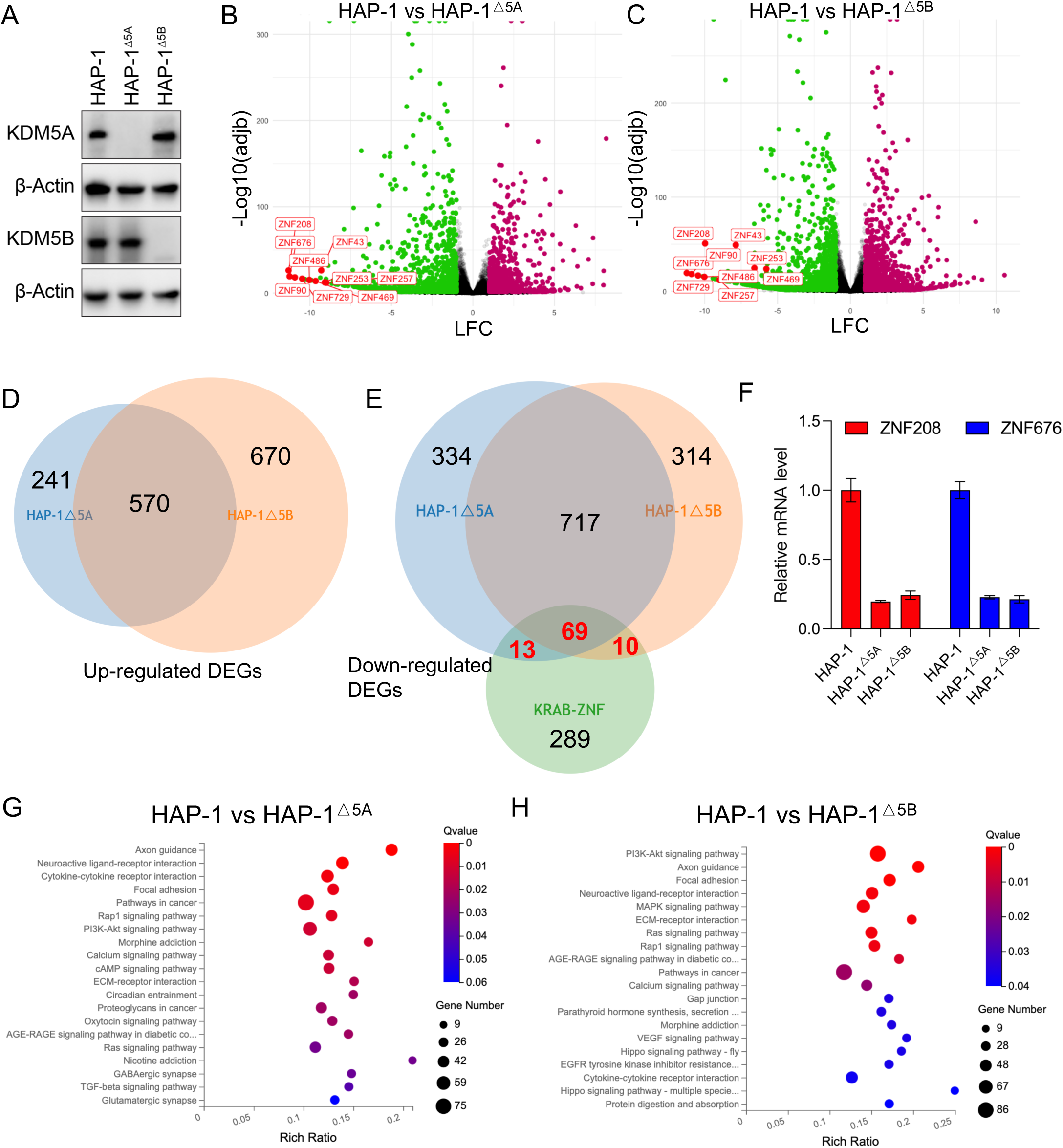
KRAB-ZNFs expression is lost in HAP1^Δ5A^ and HAP1^Δ5Β^ cells. **A** Expression of KDM5A and KDM5B assessed by western blotting of cell lysates from HAP1, HAP1^Δ5A^ and HAP1^Δ5Β^ cells. Uncropped blots are shown in Supplementary Figure S5. **B** Volcano plot showing changes in gene expression between HAP1 and HAP1^Δ5A^ cells as measured by RNA-seq. Green dots represent significantly down-regulated genes (log2FC ≤ −1, Padj ≤ 0.05), and red dots represent significantly up-regulated genes (log2FC ≥1, Padj ≤ 0.05). Red text indicates KRAB-ZNFs. **C** Volcano plot showing changes in gene expression between HAP1 and HAP1^Δ5Β^ cells as measured by RNA-seq. Green dots represent significantly down-regulated genes (log2FC ≤ −1, Padj ≤ 0.05), and magenta dots represent significantly up-regulated genes (log2FC ≥1, Padj ≤ 0.05). KRAB-ZNF genes are indicated in red text. **D** A Venn diagram showing the overlap of genes that are upregulated in HAP1^Δ5A^ and HAP1^Δ5Β^ compared to HAP1 parental cells. Upregulated genes were defined as log2FC ≥1, Padj ≤ 0.05 **E** A Venn diagram showing the overlap of genes that are downregulated in HAP1^Δ5A^ and HAP1^Δ5Β^ compared to HAP1 parental cells and the set of KRAB-ZNF genes. Downregulated genes were defined as log2FC ≤ −1, Padj ≤ 0.05. **F** RT-qPCR analyses of ZNF208 and ZNF676 mRNA levels in HAP1, HAP1^Δ5A^ and HAP1^Δ5Β^ cells. Data are shown as mean ± SEM. **G** KEGG pathway enrichment bubble charts of differentially expressed genes in HAP1^Δ5A^ compared with HAP1 parental cells. The size of the bubble represents the number of genes in each pathway, the color change represents the Qvalue, and red represents high significance. **H** KEGG pathway enrichment bubble charts of differentially expressed genes in HAP1^Δ5Β^ compared with HAP1 parental cells. The size of the bubble represents the number of genes in each pathway, the color change represents the Qvalue, and red represents high significance.

**Table 1.** Top 16 down-regulated differentially expressed genes (DEGs) between HAP1^Δ5A^ and HAP1^Δ5Β^ cells and parental cells.

| Rank | Gene | Log2(5A/WT) | Log2(5B/WT) | Relevant pathways |
| --- | --- | --- | --- | --- |
| 1 | ZNF208 | -11.309665 | -9.9724598 | KRAB-ZNF |
| 2 | ZNF676 | -11.245997 | -11.230806 | KRAB-ZNF |
| 3 | CNTNAP5 | -10.992435 | -10.977532 | Cell Adhesion |
| 4 | ZNF90 | -10.919376 | -10.904046 | KRAB-ZNF |
| 5 | RFLNA | -10.508441 | -10.492941 | Actin Filament Bundle Organization |
| 6 | ZNF729 | -10.472286 | -10.456968 | KRAB-ZNF |
| 7 | TKTL1 | -10.287032 | -8.4138657 | Glucose Catabolic Process |
| 8 | ZNF486 | -10.037081 | -10.021834 | KRAB-ZNF |
| 9 | FGF4 | -9.9196742 | -3.6687554 | Embryonic Development and Cell Proliferation |
| 10 | MLF1 | -9.8934386 | -0.6989044 | Cell Cycle Arrest |
| 11 | ZNF469 | -9.6470944 | -5.7744094 | Transcriptional Regulation |
| 12 | TMEM108 | -9.3577305 | -6.2545084 | Neuron Projection Development |
| 13 | ZNF43 | -9.2990779 | -7.8688311 | KRAB-ZNF |
| 14 | ZNF253 | -9.0812729 | -6.5849217 | KRAB-ZNF |
| 15 | ARHGEF15 | -9.0076419 | -8.4076995 | Activation Of GTPase Activity |
| 16 | ZNF257 | -9.0023291 | -8.9874562 | KRAB-ZNF |

**Table 2.**
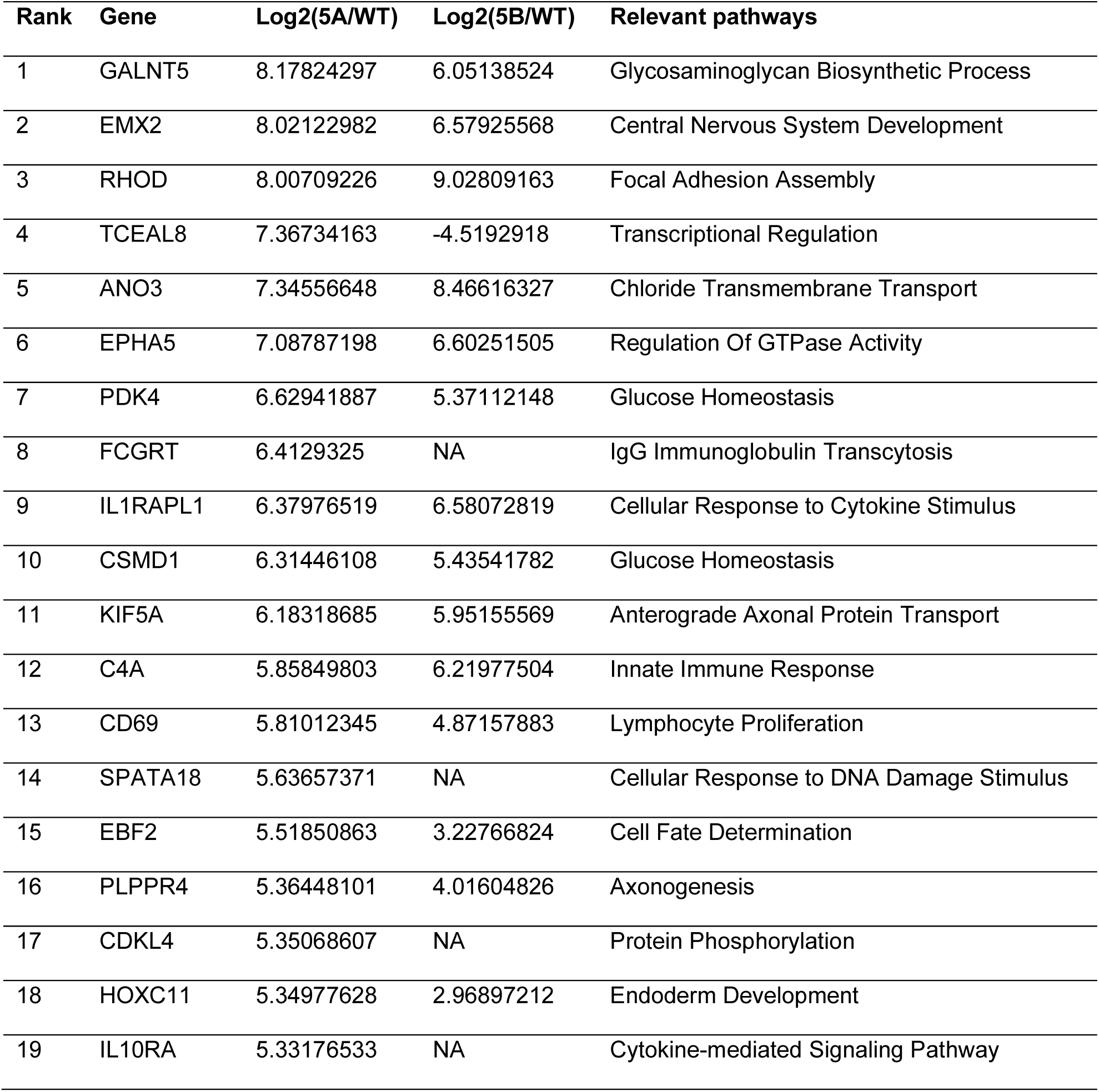
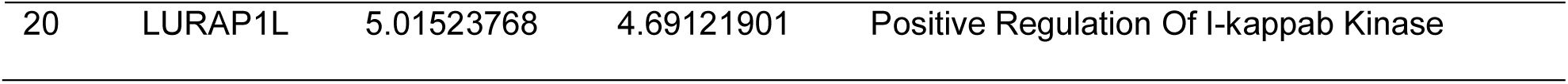
Top 20 up-regulated DEGs between HAP1^Δ5A^ and HAP1^Δ5Β^ cells and parental cells.

We next used ATAC-seq to assay global chromatin accessibility in HAP1^Δ5A^ and HAP1^Δ5Β^ cells compared to parental HAP1 cells. MACS2 peak caller was used to identify accessible regions in duplicate samples of HAP1, HAP1^Δ5A^ and HAP1^Δ5Β^. Among the genomic loci with differential accessibility between wild-type and mutant cells, multiple KRAB-ZNFs gene clusters were less open in both HAP1^Δ5A^ and HAP1^Δ5Β^ cells, consistent with decreased transcription of these targets (Fig. 2D and Table 3). While there were some variabilities in the FRIP (fraction of reads in peak) scores among the different sample groups, each sample had a high FRIP score and the replicates within each group showed largely similar peak statistics. Peak annotation analysis revealed comparable peak distributions between WT, KDM5A and KDM5B samples with the majority of peaks occurring in intronic and intergenic regions. Our analysis also found no global changes in peak density around transcriptional start sites (TSS), merged peak regions and gene bodies (Fig. S2). Despite similar global profiles, PCA analysis revealed a clear separation among the different sample groups (Fig. 2A), and pairwise comparisons between the WT, KDM5A, and KDM5B samples identified approximately 175,000 differential peaks with an adjusted p-value cutoff of < 0.1. Of these peaks, 21,271 were only present in parental cells, 12,317 (7%) were specific to KDM5A mutant cell and 7,990 (4.5%) were specific to KDM5B mutant cells. We also identified 18,548 peaks that were present in HAP1^Δ5A^ and HAP1^Δ5Β^ cells but not parental cells, 5,156 open regions that were present in HAP1^Δ5Β^ and WT cells but absent in HAP1^Δ5A^ cells and 11,933 (7%) peaks that were present in HAP1^Δ5A^ and HAP1 cells but lost in HAP1^Δ5Β^ cells and 18,548 peaks that were only present in HAP1^Δ5A^ and HAP1^Δ5Β^ cells (Fig. 2B). In addition, GO pathway analysis on genes related to chromatin accessibility in HAP1^Δ5A^ and HAP1^Δ5Β^ revealed that many olfactory receptors (OR) and G-protein coupled receptor-related genes had low chromatin accessibility (Fig. 2C). In addition, the chromatin state for many genes related to complement activation and immune response was more open in HAP1^Δ5A^ and HAP1^Δ5Β^ cells (Fig. 2C).

**Fig.2.**
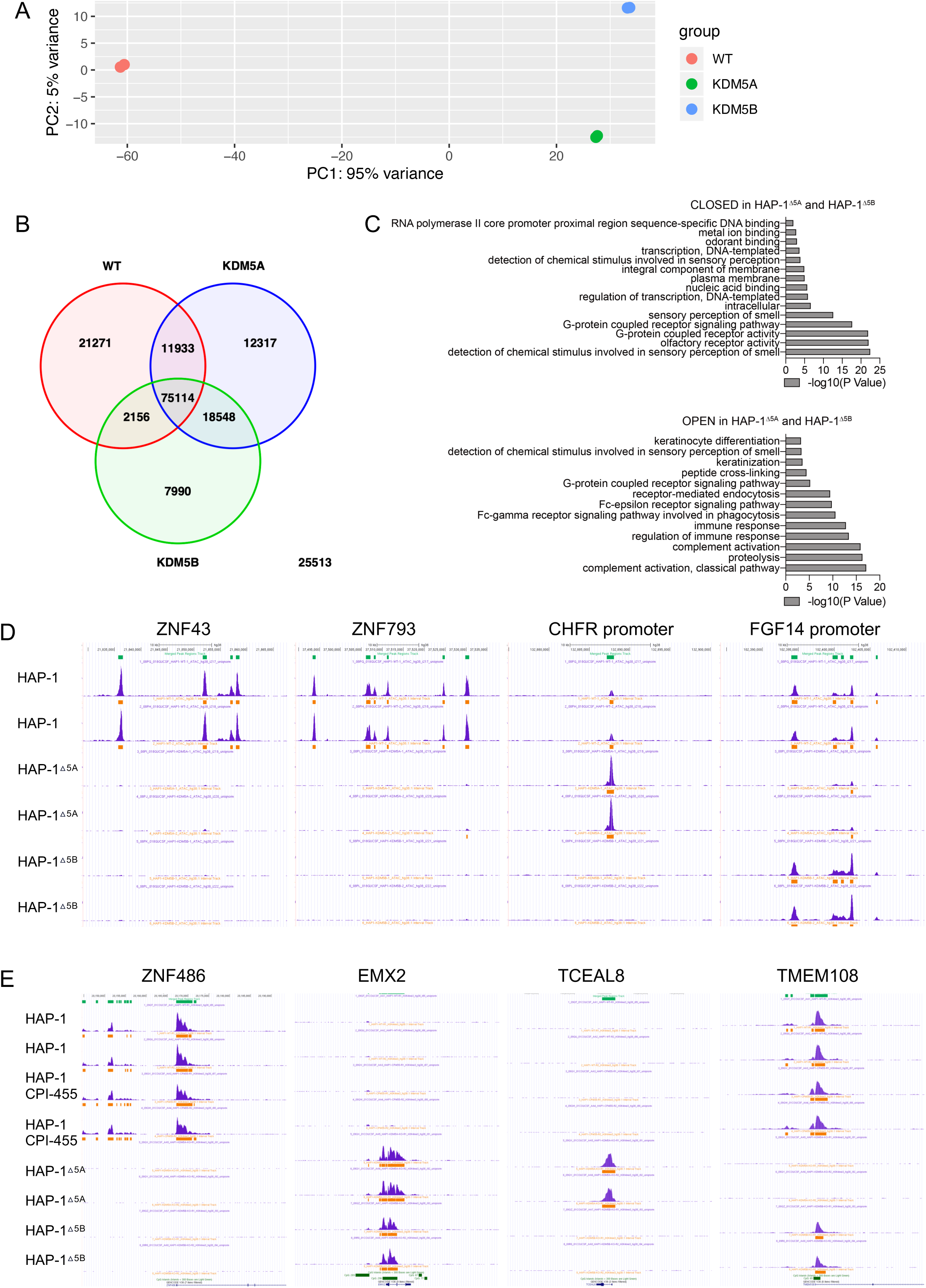
KRAB-ZNFs chromatin accessibility loss in HAP1^Δ5A^ and HAP1^Δ5Β^ cells assayed by ATAC-Seq and H3K4me3 ChIP-Seq. **A** Principal component analysis (PCA) plot of HAP1, HAP1^Δ5A^ and HAP1^Δ5Β^ ATAC-seq data. Clustering reveals greatest variance between HAP1^Δ5A^ and HAP1^Δ5Β^ cells compared to parental in PC1 with clear separation of HAP1^Δ5A^ and HAP1^Δ5Β^ cells in PC2. **B** A Venn diagram showing that overlap of ATAC-seq peaks that were present in HAP1^Δ5A^, HAP1^Δ5Β^ and wild-type cells. Only peaks that were consistent between replicates are included in this analysis; 25,513 merged regions that showed discordance between duplicate samples were excluded. **C** The top Gene Ontology (GO) terms of significantly “OPEN” or “CLOSED” genes in HAP1^Δ5A^ and HAP1^Δ5Β^ cells. **D** Normalized ATAC-seq alignments showing regional differences in chromatin accessibility surrounding ZNF43, ZNF793, CHFR promoter and FGF14 in HAP1, HAP1^Δ5A^ and HAP1^Δ5Β^ cells. The tracks were visualized using the UCSC genome browser. **E** Normalized H3K4me3 ChIP-Seq alignments showing regional differences in H3K4me3 peaks surrounding the ZNF486, EMX2, TCEAL8 and TMEM108 loci in HAP1, HAP1^Δ5A^ and HAP1^Δ5Β^ and CPI-455 treated HAP1. The tracks were visualized using the UCSC genome browser.

**Table 3.** ATAC chromosome accessibility status of top 16 KRAB-ZNFs between HAP1^Δ5A^ and HAP1^Δ5Β^ and parental cells.

| Rank | Gene | Chromatin Access | Chr |
| --- | --- | --- | --- |
| 1 | ZNF208 | Closed in KO | chr19 |
| 2 | ZNF676 | Closed in KO | chr19 |
| 3 | ZNF90 | Closed in KO | chr19 |
| 4 | ZNF729 | Closed in KO | chr19 |
| 5 | ZNF486 | Closed in KO | chr19 |
| 6 | ZNF43 | Closed in KO | chr19 |
| 7 | ZNF257 | Closed in KO | chr19 |
| 8 | ZNF626 | Closed in KO | chr19 |
| 9 | ZNF99 | Closed in KO | chr19 |
| 10 | ZNF667 | Closed in KO | chr19 |
| 11 | ZNF264 | Closed in KO | chr19 |
| 12 | ZNF433 | Closed in KO | chr19 |
| 13 | ZNF844 | Closed in KO | chr19 |
| 14 | ZNF578 | Closed in KO | chr19 |
| 15 | ZNF586 | Closed in KO | chr19 |
| 16 | ZNF98 | Closed in KO | chr19 |

As KDM5A and KDM5B regulate the methylation status of histone H3 lysine 4 (H3K4), we conducted ChIP-Seq analysis to assess the genomic distribution of H3K4me3 modifications in HAP1^Δ5A^ and HAP1^Δ5Β^ cells as well as HAP1 cells treated with the potent and selective KDM5 demethylase inhibitor, CPI-455. Although the overall distribution of H3K4me3 was consistent across all samples, principal component analysis (PCA) revealed a clear separation between HAP1^Δ5A^ and HAP1^Δ5Β^ cells compared to parental HAP1 cells. In contrast, CPI-455 treated HAP1 cells did not show significant separation from parental cells (Fig. S3A). Similar to our ATAC-Seq results, ChIP-Seq analysis showed no global alterations in peak density surrounding transcriptional start sites (TSS), merged peak regions, and gene bodies (Fig. S3B) in HAP1^Δ5A^ and HAP1^Δ5Β^ cells and CPI-455 treated HAP1 cells compared to parental HAP1 cells. However, we found that the H3K4me3 peak at KRAB-ZNFs genes completely disappeared in HAP1^Δ5A^ and HAP1^Δ5Β^ cells compared to HAP1 parental cells, consistent with the results from RNA-Seq and ATAC-Seq. In contrast, CPI-455 did not alter H3K4me3 within the KRAB-ZNFs loci demonstrating that inhibition of enzymatic activity was not sufficient to alter the chromatin in these regions (Fig. 2E). Taken together, these results indicate that both KDM5A and KDM5B are required to maintain open chromatin and transcriptional activity of KRAB-ZNFs gene clusters. Furthermore, the regulation of KRAB-ZNF by KDM5A and KDM5B appears independent of their catalytic activity, as evidenced by the lack of an effect of CPI-455 in the ChIP-Seq analysis.

### Enhanced ERV transcription and immune response in KDM5A/B knockout cells

KRAB-ZFPs comprise the largest family of transcriptional repressors in the human genome [12]. Approximately two thirds of the human genome consist of transposable elements which are, in part, transcriptionally repressed by KRAB-ZFPs [13,14]. Ablation of *KDM5B* is associated with upregulation of ERVs, including ERV-T and ERV-S during development. These observation led us to hypothesize that decreased expression of repressive KRAB-ZNFs in HAP1^Δ5A^ and HAP1^Δ5Β^ cells may lead to an increase in ERV transcription [10]. Owing to the high homology between and within proviral loci, quantification of ERV transcripts is complicated because sequencing reads from ERV RNAs often align with multiple loci. Additionally, HERV-K proviruses are poorly annotated in human transcriptome databases making their analysis in RNA-seq data difficult [15,16]. Therefore, to investigate changes in RNA levels of ERVs we used RT-qPCR to measure steady state levels of select ERV transcripts in parental and knockout lines cells. These experiments showed that HAP1^Δ5A^ and HAP1^Δ5Β^ cells had increased levels of *ERV3-1*, *ERVW*, *HERVE* and *HERVF* transcripts (Fig. 3A). Inhibition of KDM5A and KDM5B catalytic activity with the small molecule inhibitor CPI-455 did not alter levels of these transcripts in HAP1, HAP1^Δ5A^ or HAP1^Δ5Β^ cells, suggesting that suppression of ERV transcription by KDM5A and KDM5B is independent of their catalytic activity (Fig. S4A). Increased transcription of ERVs can lead to an accumulation of cytoplasmic dsRNA and trigger an immune response [17–19]. To explore this possibility, we performed immunofluorescence analysis to measure the levels of dsRNA in HAP1, HAP1^Δ5A^ and HAP1^Δ5Β^ cells and found that HAP1^Δ5A^ and HAP1^Δ5Β^ have elevated levels of dsRNA foci in the cytoplasm compared to HAP1 parental cells. Consistent with our observation that CPI-455 did not increase *ERV* transcripts, CPI-455 treatment did not affect the number of dsRNA foci (Fig. 3B and 3C). Loss of KDM5A or KDM5B was also associated with an increase in *ERV* transcripts in a chronic myelogenous leukemia (CML) derived K562 cell line (Fig. 3D and 3E); however, we found no evidence of elevated levels of dsRNA in K562^Δ5A^ and K562^Δ5Β^ cells (Fig. S4C). As in HAP1 cells, CPI-455 had a modest to no effect on ERV mRNA levels in K562, K562^Δ5A^ or K562^Δ5Β^ cells (Fig. S4B).

**Fig.3.**
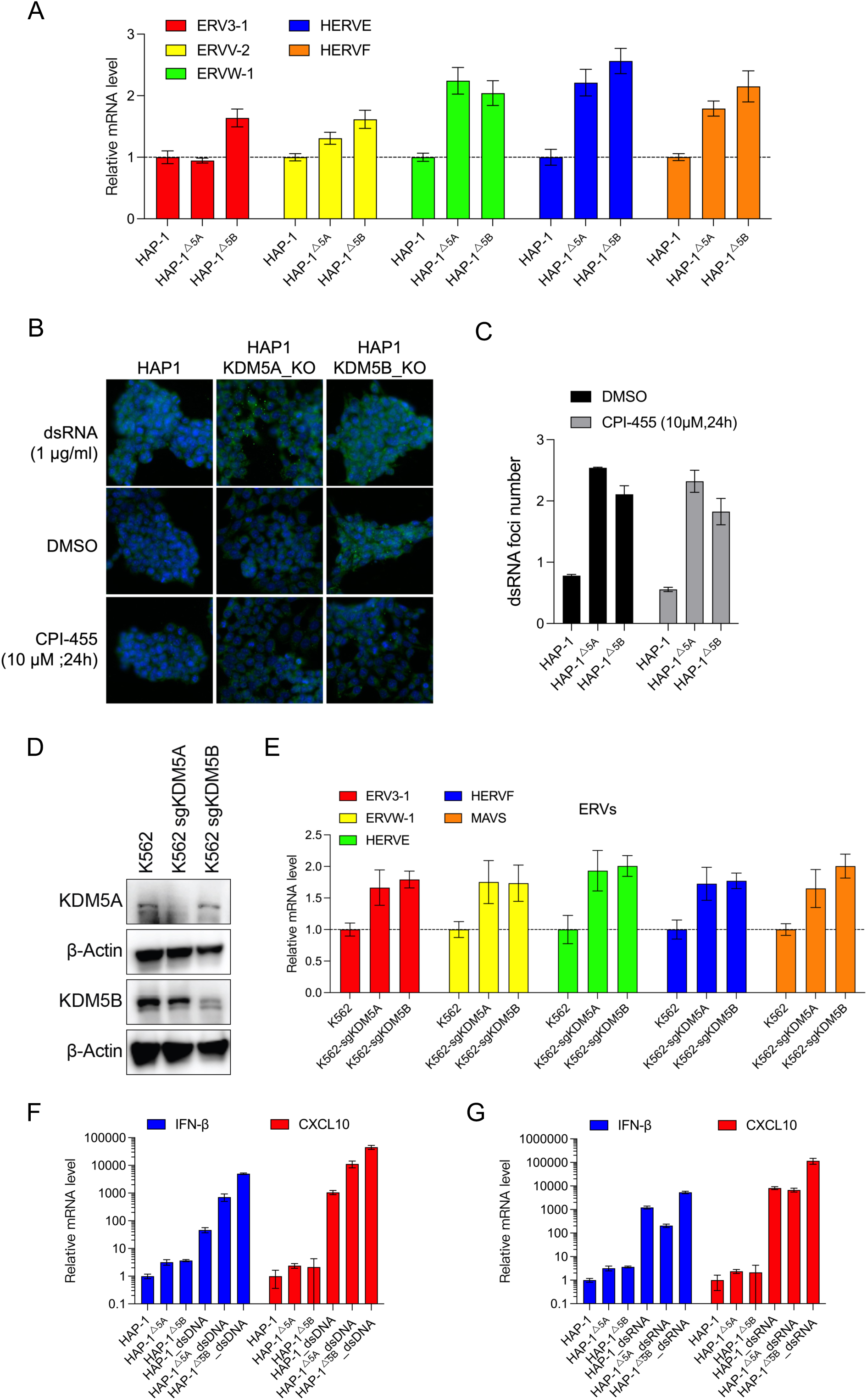
Enhanced transcription of ERVs and immune response in KDM5A/B knockout cells. **A** RT-qPCR analyses of ERVs gene transcription levels in HAP1^Δ5A^ and HAP1^Δ5Β^ cells and HAP1 cells. Data are mean ± SEM. **B** Representative images of dsRNA levels in HAP1^Δ5A^ and HAP1^Δ5Β^ cells, HAP1 cells, or CPI-455 treated HAP1 cells. Cells were stained with dsRNA Rabbit mAb and Goat anti-Rabbit IgG Alexa Fluor 488 secondary antibody. DMSO treated cells are shown as a vehicle control. **C** Quantification of dsRNA Alexa 488 cytoplasmic foci from cells in (**b**). Data shown as mean values ± SD; At least 10,000 cells were analyzed in each group, from triplicate wells. **D** Expression of KDM5A and KDM5B assessed by western blotting of cell lysates from K562, K562^Δ5A^ and K562^Δ5Β^ cells. β-actin levels shown as a loading control. Uncropped blots are shown in Supplementary Figure S6. **E** RT-qPCR analyses of ERV transcript levels in K562, K562^Δ5A^ or K562^Δ5Β^ cells. Data are plotted as mean ± SEM. **F** RT-qPCR analyses of IFN-β and CXCL10 transcript levels in HAP1, HAP1^Δ5A^ and HAP1^Δ5Β^ cells incubated with and without dsDNA. Data are mean ± SEM. **G** RT-qPCR analyses of IFN-β and CXCL10 transcript levels in HAP1, HAP1^Δ5A^, HAP1^Δ5Β^ cells incubated with and without dsRNA. Data are mean ± SEM.

In both HAP1^Δ5A^ and HAP1^Δ5Β^ cells, the increased number of dsRNA foci was associated with an increase in IFN-β and CXCL10 mRNA levels (Fig. 3F and 3G), suggesting that loss of epigenetic regulation by KDM5A and KDM5B can enhance immune signaling. To further explore this possibility, we treated HAP1, HAP1^Δ5A^ and HAP1^Δ5Β^ cells with exogenous dsRNA and dsDNA. This experiment revealed that dsDNA and dsRNA both caused an increase in steady state levels of IFN-β and CXCL10 transcripts across all three cell lines, with the most dramatic increase in mRNA levels occurring in HAP1^Δ5Β^ cells treated with dsRNA (Fig. 3F and 3G). This difference supports the conclusion that KDM5A and KDM5B have both overlapping and distinct functions. Again, CPI-455 treatment did not alter how HAP1 cells respond to dsDNA and dsRNA (Fig. S4D), suggesting that the upregulation of immune-related transcripts is independent of KDM5A and KDM5B enzymatic activity.

### dTAG-mediated degradation of KDM5A stimulates ERV expression

To characterize the role of KDM5A in suppressing ERV expression, we used a dTAG approach to chemically induce degradation of KDM5A protein. The dTAG system utilizes a heterobifunctional small molecule that specifically binds and brings in close proximity a FKBP12^F36V^-tagged protein and the E3 ligase complex, leading to ubiquitination and proteasome-mediated degradation of the target protein [20,21]. To engineer a cell model in which we could conditionally degrade KDM5A, we used CRISPR-Cas9 to knock-in an FKBP12^F36V^-2xHA tag to the N-terminus of the sole copy of KDM5A in the haploid HAP1 cells (Fig. 4A). This ‘dTAG-KDM5A’ fusion protein can be degraded by a bifunctional degrader, dTAG-47, which comprises ligands specific for FKBP12^F36V^ and the E3 ligase cereblon which targets the modified KDM5A for cereblon-mediated ubiquitination and subsequent proteolytic degradation [22]. dTAG-47 can degrade the dTAG-KDM5A chimera in a dose-dependent manner, with optimal degradation observed at a concentration of 0.5 µM, resulting in the removal of >90% of the fusion protein (Fig. 4B). The characteristic hook effect behavior, which is commonly observed with heterobifunctional degraders due to saturation of FKBP12^F36V^ and E3 ligase binding sites [23] was evident at higher dTAG-47 concentration (5 µM). Time-course experiments revealed fast kinetics of dTAG-KDM5A degradation with >90% of the fusion protein being degraded within four hours (Fig. 4C). Next, we assessed the effect of dTAG-47-induced KDM5A loss on *ERV* gene expression. We performed RT-qPCR for select human *ERV* genes from mRNA collected from dTAG-47 treated HAP1^dTAG-KDM5A^ cells (Fig. 4D). Acute loss of KDM5A led to increased transcript levels of *ERV3-1*, *ERVV-2*, *ERVW*, and *HERVE*, whereas changes in *HERVF* transcript levels were insignificant. Next, we measured how acute loss of KDM5A affected steady state levels of two of the most downregulated transcripts identified by RNA-seq in HAP1^Δ5A^ cells, ZNF208 and ZNF676. Surprisingly, no significant changes were observed in the levels of *ZNF208* or *ZNF676* transcripts following dTAG-47 treatment (Fig. 4D). The different effects of genetic ablation and dTAG-induced loss of KDM5A on ZNF expression could be attributed to the knockout cells adapting to KDM5A deficiency.

**Fig.4.**
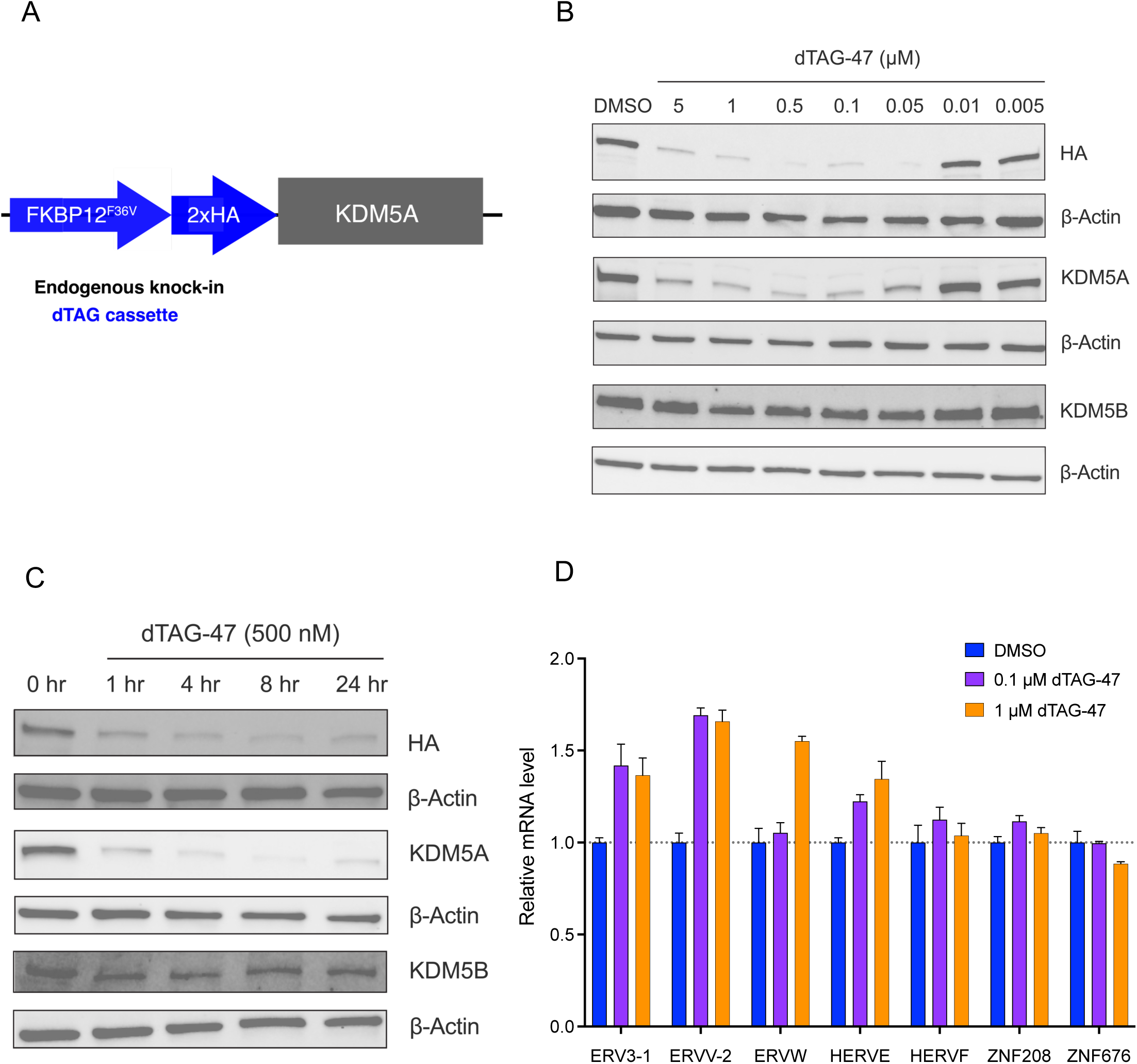
Acute loss of KDM5A using dTAG depletion induces ERV expression. **A** Schematic depiction of dTAG knock-in onto the N-terminus of KDM5A in HAP1 cell line. The dTAG cassette, comprising of FKBP12^F36V^ and 2X HA-tag, was inserted into the KDM5A locus using CRISPR-Cas9. **B** Immunoblot analysis of HAP1^dTAG-KDM5A^ cells treated with control (DMSO) or dTAG-47 at the indicated doses for 4 hours. Uncropped blots are shown in Supplementary Figure S7. **C** Kinetics of dTAG-KDM5A degradation in HAP1^dTAG-KDM5A^ cells following dTAG-47 treatment (500 nM). Uncropped blots are shown in Supplementary Figure S8. **d** RT-qPCR analysis of the relative mRNA levels for select ERVs and ZNFs in HAP1^dTAG-KDM5A^ cells treated with DMSO or dTAG-47 (0.1 µM or 1 µM) for 48 hours. Data are mean ± SEM.

### KDM5A is a part of KRAB-ZNF repressor complex

KRAB-ZNFs are known to bind KAP1 (also known as TRIM-28), a transcriptional repressor that interacts with the KRAB repression domain found in many transcription factors [24,25]. KAP1 serves as a scaffolding protein for recruitment of chromatin-related corepressors: SETDB1, a histone H3K9me3 methyltransferase and the NuRD complex, responsible for deacetylation of lysine on histone tails and nucleosome remodeling [26,27]. It has been shown previously that, in HeLa cells, KDM5A associates with HDAC1/2, histone-binding protein RBAP46/48, ATP-dependent chromatin remodeler (CHD3/CDH4), metastasis-associated factor (MTA1/MTA2/MTA3), methyl-DNA-binding protein (MBD2/MBD3), and GATAD2 [28,29], which are the components of NuRD complexes. To evaluate if KDM5A associates with these silencing complexes in HAP1 cells, we performed co-immunoprecipitation (co-IP) and subsequent immunoblotting for components of both complexes (Fig. 5A). Immunoprecipitation of a HA-tagged KDM5A pulled down two protein components of the KRAB-ZNF repressor complex, KAP1 and SETDB1. Immunoprecipitation also revealed an interaction between KDM5A and components of NuRD: RBAP46, HDAC1, HDAC2, MTA1 and MBD3. These results provide evidence that KDM5A interacts with both the KAP1-SETDB1 repressor complex and the NuRD complex. These findings highlight a novel interaction between KDM5A, SETDB1 and KAP1, suggesting the potential role of KDM5A as a component of KRAB-ZNF repressor complex.

**Fig. 5.**
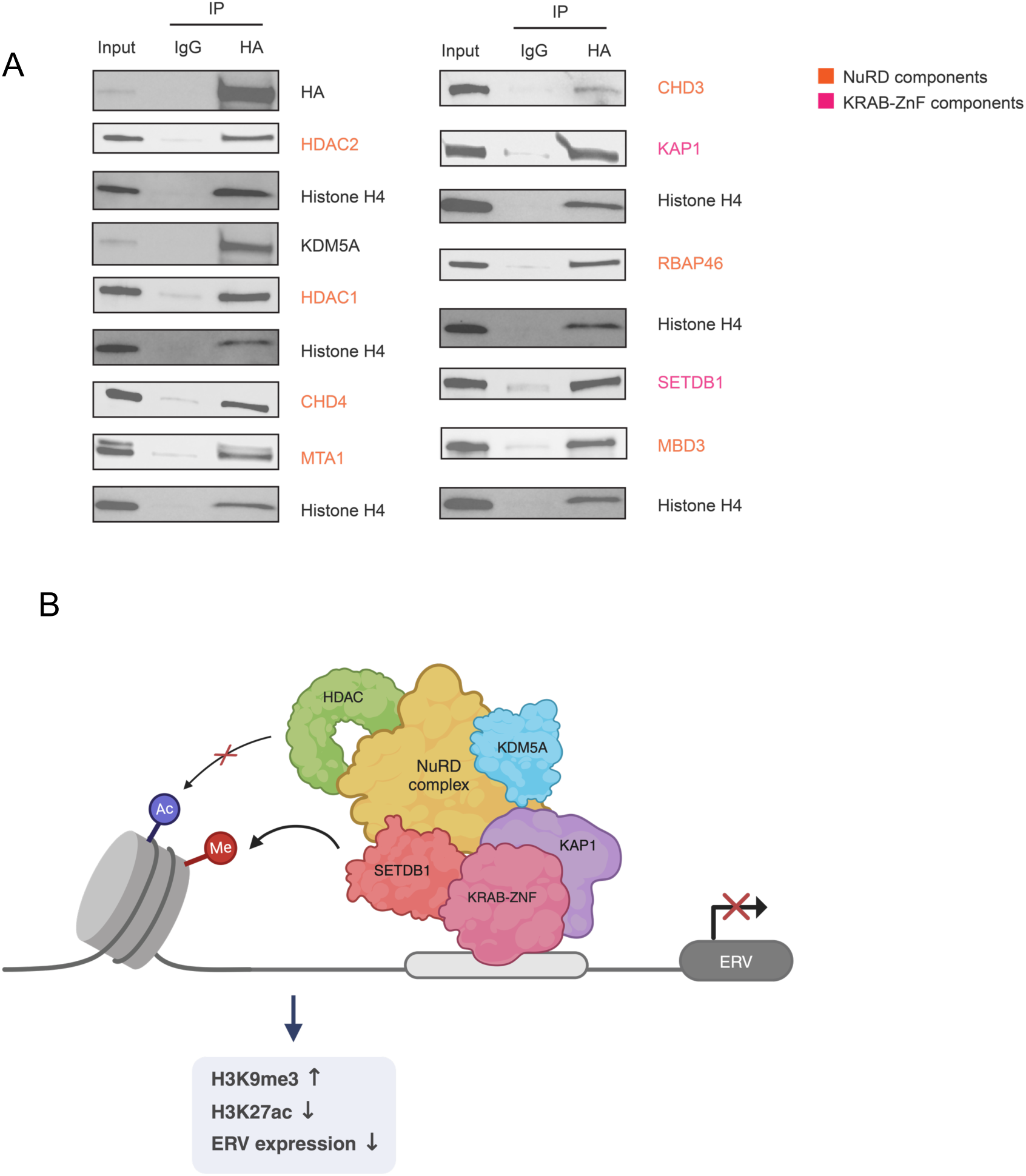
KDM5A associates with KRAB-ZNF and NuRD components. **A** Co-immunoprecipitation of dTAG-KDM5A from HAP1^dTAG-KDM5A^ cell extracts, followed by immunoblotting for subunits of NuRD and KRAB-ZNF complexes. Western blots were performed on multiple gels, with histone H4 as a loading control for each gel. Uncropped blots are shown in Supplementary Figure S9. **B** A model for suppression of ERV gene expression. KDM5A may facilitate the recruitment of SETDB1 and NuRD components to the KAP1-KRAB-ZNF repressor complex to enforce silencing of ERVs. Figure created with Biorender.com.

## Discussion

The KDM5 proteins, members of the Jumonji C (JmjC) domain-containing histone demethylase family, play crucial roles in epigenetic regulation by catalyzing the demethylation of histone H3 lysine 4 (H3K4) [30,31]. These enzymes are notable for their involvement in diverse biological processes, including transcriptional regulation and cell differentiation [32]. Moreover, dysregulation of KDM5A and KDM5B enzymes has been implicated in various human diseases, particularly cancer, underscoring their potential as therapeutic targets [33]. Through their scaffolding functions, KDM5A and KDM5B can mediate interactions with various transcription factors and chromatin-modifying complexes [34–37]. Recent studies have highlighted the significance of KDM5A and KDM5B in regulating immune checkpoints, cytokine production, and the inflammatory response [8–10]; dysregulation of KDM5A and KDM5B expression or activity has been implicated in the pathogenesis of autoimmune diseases, inflammatory disorders, and cancer immune evasion [38]. Understanding the roles of KDM5A and KDM5B in immune regulation holds promise for developing novel immunotherapeutic strategies and targeted interventions for cancer and immune-related diseases.

Here, we employed a multi-omics approach to investigate the roles of KDM5A and KDM5B in chromatin regulation. Global RNA-seq analysis revealed multiple pathways affected by loss of KDM5A and KDM5B (Fig. 1). Notably, deletion of KDM5A or KDM5B in HAP1 cells resulted in transcriptional repression of *KRAB-ZNF* genes. This observation was further validated by ATAC-seq and H3K4me3 ChIP-seq experiments, which showed low chromatin accessibility and H3K4me3 loss at *KRAB-ZNF* loci in both KDM5A and KDM5B knockout HAP1 cells (Fig. 2). KRAB-ZNFs are known to suppress ERV transcription [12] and our data reveal that decreased expression of KRAB-ZNFs in HAP1^Δ5A^ and HAP1^Δ5Β^ cells is correlated with higher expression of ERV transcripts, increased levels of dsRNA and elevated levels of immune response genes, including IFN-β and CLXCL10. While similar effects were seen in another CML cell line, K562, after knockout of KDM5A or KDM5B (Fig. 3E), these phenotypes were not present in other cancer cell lines that we tested. Additionally, the role of KDM5B in regulating ERV expression has also been noted in mouse and human melanoma cell lines. A previous report has shown coregulation of ERV expression by KDM5B and SETDB1 in these cell lines, suggesting that distinct mechanisms may be employed to repress ERV expression in different genetic contexts [38].

Small molecule inhibitors of KDM5 have been developed in recent years to suppress KDM5 activity across various disease models [39–43]. Amongst these inhibitors, GS-5801, an oral liver-targeted KDM5 inhibitor for Hepatitis B, currently remains the only inhibitor to have reached the clinic [44]. Unfortunately, GS-5801 failed in an early phase clinical trial due to tolerability concerns [45], highlighting the ongoing challenge and unmet needs in KDM5 therapy. The multifaceted roles of KDM5A and KDM5B also complicate the development of small molecule inhibitors, as blocking enzymatic activity may not affect non-catalytic functions that contribute to disease progression. Currently, all KDM5 small molecule inhibitors target the catalytic site of the protein. In this study, we used CPI-455, a pan-KDM5 orthosteric inhibitor, previously shown to increase global level of H3K4me3 and decrease the number of drug tolerant population in multiple cancer cell line models. Surprisingly, treating cells with CPI-455 did not result in the downregulation of KRAB-ZNF genes or alter the H3K4me3 status of *KRAB-ZNF* gene targets, suggesting that regulation of these targets occurs independently of demethylase activity (Fig S4A, S4B). Several recent studies point to an emerging role of the scaffolding function of KDM5 proteins in regulation of gene expression. In a mouse melanoma model, KDM5B is reported to recruit a H3K9me3 methyltransferase, SETDB1, to promote immune evasion through silencing of transposable elements, independently of KDM5B demethylase activity [10]. The catalysis-independent function of KDM5 proteins extends beyond cancer models and has been observed in *Drosophila*, where a recruitment function through a chromatin reader domain in KDM5 is essential for the regulation of gene expression [46]. We note that loss of either KDM5A or KDM5B also resulted in lower KRAB-ZNF expression and increased ERV expression, indicating that the functions of KDM5A and KDM5B in regulating KRAB-ZNF are not redundant.

To probe if additional mechanisms may contribute to the ability of KDM5A to downregulate *ERV* expression, we investigated its association with known repressive factors. A previous study reported association of KDM5A and KDM5B with components of the NuRD complex, where they cooperatively function to control developmentally regulated genes [28,47]. NuRD has also been shown to interact with the KRAB-ZFP repressor complex to deacetylate histones in the promoter regions for effective gene silencing [48]. Through co-immunoprecipitation, we found that KDM5A associates with the components of NuRD complex and KRAB-ZFP complex. These results provide evidence that, by facilitating protein-protein interactions, KDM5A cooperates with NuRD, KAP1, and SETDB1 to enforce silencing of ERVs (Fig. 5B). The inability of KDM5A inhibitors to cause reactivation of *ERV* genes supports a demethylase-independent function. Future studies are warranted to explore whether targeting the interactions within the repressive complex can be leveraged to reactivate ERVs.

Proteolysis-targeting chimeras (PROTACs) are a rapid and selective method for reducing the abundance of target proteins, abolishing both the catalytic and non-catalytic functions such as scaffolding [49,50]. Compared with traditional gene-editing approaches, acute removal of targets using PROTACs can provide insights on the direct effects of degradation without those effects being confounded by adaptation or secondary effects [51]. To investigate the mechanisms by which KDM5A proteins contribute to transcriptional repression of *ERV* genes, we utilized the dTAG system to chemically induce degradation of KDM5A [20,21]. Immediately following the loss of KDM5A, we observed an increase in *ERV* expression, consistent with the effects seen in HAP1^Δ5A^ and HAP1^Δ5Β^ models. The timing of this response suggests that de-repression of *ERV* elements is an acute response to KDM5A loss and not an adaptive change that occurs in cells that are deficient for KDM5A. Notably, there was no significant alteration in *ZNF208* and *ZNF676* transcript levels after acute KDM5A loss. It is possible that sustained decreases in these ZNF transcripts may not be achievable within the experimental timeframe, suggesting that KDM5A may influence *ERV* expression through both ZNF-dependent and ZNF-independent pathways.

In summary, we have generated a rich dataset for the exploration of KDM5A and KDM5B function. Our multiomics analyses identified KDM5A and KDM5B as regulators of KRAB-ZNFs. In addition, we show that genetic deletion of *KDM5A* and *KDM5B* or protein degradation of KDM5A induces ERV expression and causes an enhanced immune response characterized by increased dsRNA and elevated expression of CXCL10 and IFN-β. This immune-suppressive activity of KDM5A and KDM5B is independent of demethylase activity, adding to a growing repertoire of data supporting the crucial scaffolding function of these enzymes and providing further support for developing KDM5A and KDM5B degraders as immune modulatory anti-cancer treatments.

## Materials and methods

### Cell culture

Human K562 cells were obtained from the American Type Culture Collection (ATCC, Manassas VA, USA). Human HAP1 (C631) and HAP1^Δ5A^ (HZGHC004366c001) and HAP1^Δ5Β^ (HZGHC004164c008) cells were purchased from Horizon Discovery. K562 cells were grown in Roswell Park Memorial Institute (RPMI) 1640 Medium (ATCC modification) with 10% fetal bovine serum and 1% penicillin-streptomycin. HAP1 cells were grown in Iscove’s Modified Dulbecco’s Medium (IMDM) with 10% fetal bovine serum and 1% penicillin-streptomycin. HAP1 and K562 cells were authenticated by short tandem repeat (STR) profiling and tested for mycoplasma at Genetica.

### Construction of plasmids

The lentiCRISPRv2 sgRNA plasmids were constructed using the method previously described by the Zhang lab (9,10) and the sgRNA targeting sequences used are as follows: KDM5A-targeting sgRNA (5′-GTGTCCTAAATGTGTCGCCG-3′) and KDM5B-targeting sgRNA (5′-TCTTGCAGATCATCTCATCG-3′). A detailed protocol is available at https://media.addgene.org/cms/filer_public/4f/ab/4fabc269-56e2-4ba5-92bd-09dc89c1e862/zhang_lenticrisprv2_and_lentiguide_oligo_cloning_protocol_1.pdf.

### RNA seq

Five million HAP1, HAP1^Δ5A^ and HAP1^Δ5Β^ cells in the exponential proliferation were collected respectively. RNA was isolated using RNeasy Mini Kit (QIAGEN, Cat# 74104) and treated with DNase (QIAGEN Cat# 79254) to remove genomic DNA. RNAs were then sent to BGI for RNA quality control (via Bioanalyzer), library preparation, and next-generation sequencing on an Illumina NovaSeq instrument as a fee-for-service.

### ATAC seq

One million HAP1, HAP1^Δ5A^ and HAP1^Δ5Β^ cells in the exponential proliferation were collected respectively. Chromatin preparation and sonication, transposase reaction, library amplification, and next-generation sequencing on an Illumina NovaSeq instrument was performed by Active Motif.

### H3K4me3 ChIP seq

One million HAP1, CPI-455 treated HAP1, HAP1^Δ5A^ and HAP1^Δ5Β^ cells in the exponential proliferation were collected respectively. ChIP with a ChIP-validated H3K4me3 antibody, ChIP-Seq library preparation, and next-generation sequencing on an Illumina NovaSeq instrument was performed by Active Motif.

### Lentiviral packaging

Lentivirus was prepared as previously described [52]. Briefly, 15 million HEK293T cells were transfected 15 million HEK293T cells were grown overnight on 15 cm poly-L-Lysine coated dishes and then transfected with 6 ug pMD2.G (Addgene plasmid # 12259; http://n2t.net/addgene:12259; RRID:Addgene_12259), 18 ug dR8.91 (since replaced by second generation compatible pCMV-dR8.2, Addgene plasmid #8455) and 24 ug lentiCRISPR-V2 sgRNA plasmids using the lipofectamine 3000 transfection reagent per the manufacturer’s protocol (Thermo Fisher Scientific, Cat #L3000001). pMD2.G and dR8.91 were a gift from Didier Trono. The following day, media was refreshed with the addition of viral boost reagent at 500x as per the manufacturer’s protocol (Alstem, Cat #VB100). Viral supernatant was collected 48 hours post transfection and spun down at 300 g for 10 minutes, to remove cell debris. To concentrate the lentiviral particles, Alstem precipitation solution (Alstem, Cat #VC100) was added, mixed, and refrigerated at 4°C overnight. The virus was then concentrated by centrifugation at 1500 g for 30 minutes, at 4°C. Finally, each lentiviral pellet was resuspended at 100x of original volume in cold DMEM+10%FBS+1% penicillin-streptomycin and stored until use at −80°C.

### Establishment of Individual CRISPR Knockout Cells

To generate knockout clones for individual genes, K562 cells were infected with lentiCRISPR-V2 lentivirus containing sgRNAs of KDM5A or KDM5B. Infected cells were selected for 3 days with 2 μg/ml puromycin. Knockout efficiency were validated by western blotting.

### Real-time quantitative PCR

Total RNA was isolated using RNeasy Mini Kit (Qiagen, 74104), and 500 ng of total RNA was used to prepare cDNA using the PrimeScript™ RT Master Mix (TAKARA, RR036A) according to the manufacturer’s instructions. qRT-PCR was performed in triplicate for each target sequence using iTaq Universal SYBR Green Supermix (BIO-RAD, 1725121) on a Bio-Rad CFX96 using the primers in **Supplementary Table 1**.

### dsRNA subcellular distribution assay

To assess endogenous dsRNA localization, HAP1, HAP1^Δ5A^ and HAP1^Δ5Β^ cells were seeded in 96-well plate (5000 cells/well) and treated the next day with the indicated concentrations of dsRNA or CPI-455. After 24 hours of exposure to drugs, treated cells were fixed in pre-cooled methanol at −20°C for 20 min, blocked in 3% bovine serum albumin for 15 min, incubated with Anti-dsRNA-Rabbit (Millipore, MABE1134) antibodies for 1 h, and then incubated with Goat anti-Rabbit IgG Secondary Antibody, Alexa Fluor 488 (ThermoFisher, A-11008) secondary antibodies for 30 min. Final staining with DAPI for 10 minutes. Fluorescent cells were scanned by IN Cell Analyzer 6500 System and then analyzed by IN Cart (Cytiva).

### Engineering of dTAG cell line

CRISPR-Cas9 mediated knock-in cell clone of HAP1, HAP1^Δ5A^ and HAP1^Δ5Β^ were generated by Synthego Corporation (Redwood City, CA, USA). To generate these cells, Ribonucleoproteins containing the Cas9 protein and synthetic chemically modified sgRNA (sequence: 5′-CCCCACGCCCGCCAUUGCAA-3′) were electroporated into the cells to insert FKBP12F36V-2×HA-linker cassette into the N-terminus of KDM5A. Editing efficiency was assessed upon recovery, 48 hours post electroporation. Genomic DNA was extracted from a portion of the cells, PCR amplified and sequenced using Sanger sequencing. To create monoclonal cell populations, edited cell pools were seeded at 1 cell/well using a single cell printer into 96 or 384 well plates. All wells were imaged every 3 days to ensure expansion from a single-cell clone. Clonal populations are screened and identified using the PCR-Sanger genotyping strategy.

PCR reactions were performed using the following primers:

GAAATGCTGGAAAGGCTACTTG (ExtF),

CAACATTTCCTTCCACCTCCACT (ExtR) and

CAATGGGAGTGCAGGTGGAAACCATCTCCC (IntF),

CCCGCGCCTCCACTGCCACCAGATCCGCCT (IntR).

### Immunoblotting

Cells were lysed using RIPA buffer (Thermo Scientific, no. 89900) supplemented with 5X Halt Protease and Proteinase Inhibitor Cocktail with 0.5 mM EDTA (Thermo Scientific, no. 78440) and 25 U/mL Benzonase Nuclease (Millipore Sigma, no. 70746). Lysates were incubated at 4 °C on an end-over-end rocker for 30 minutes and cleared by 14,000xg centrifugation for 20 minutes at 4 °C. The total protein concentration was then measured with Bradford assay (Bio-Rad, no. 5000006). Equal amounts of protein were separated by sodium dodecyl sulfate polyacrylamide gel electrophoresis (SDS-PAGE) and transferred to NC membranes (Bio-Rad, no. 1704158). 4× Laemmli Sample Buffer (Bio-Rad, no. 1610747) supplemented with 10% β-mercaptoethanol (Millipore Sigma, no. 63689) was mixed with an equal concentration of cell lysates and boiled for 10 minutes. Samples were then loaded onto 4-20% SDS page gels (Bio-Rad, no. 4561093). Membranes were blocked in 5% non-fat milk in Tris-buffered saline (50 mM Tris-HCl, 138 mM NaCl, 2.7 mM KCl, pH 7.4) with 0.1% Tween-20 (TBS-T) and incubated with primary antibodies in the same buffer or in 5% BSA in TBS-T overnight at 4 °C. Membranes were then incubated with secondary anti-rabbit or anti-mouse antibodies for 1 h at room temperature and developed using Amersham ECL Prime Western Blotting Detection Reagent (Cytiva, no. RPN2232) and imaged using the ChemiDoc imaging system (Bio-Rad). The following antibodies were used in this study: Goat anti-rabbit IgG, HRP-linked (Cell Signaling, no. 7074, 1:3000), Horse anti-mouse IgG, HRP-linked (Cell Signaling, no. 7076, 1:3000), anti-KDM5A (Abcam, ab194286, 1:5000), anti-KDM5B (Cell Signaling, no. 3723, 1:1000), anti-HA (Cell Signaling, no. 3724, 1:1000), Anti-β-Actin (HRP conjugate; CST, 5125S) or anti-β-Actin (Cell Signaling, no. 3700, 1:2000), anti-KAP1 (Proteintech, no. 15202-1-AP, 1:1000), anti-SETDB1 (Proteintech, no. 11231-1-AP), anti-CHD3 (Cell Signaling, no. 4241, 1:1000), anti-CHD4 (Cell Signaling, no. 11912, 1:1000), anti-HDAC1 (Cell Signaling, no. 5356, 1:1000), anti-HDAC2 (Cell Signaling, no. 5113, 1:1000), anti-MBD3 (Cell Signaling, no. 14540, 1:1000), anti-MTA1 (Cell Signaling, no. 5647, 1:1000), anti-RBAP46 (Cell Signaling, no. 6882, 1:1000), anti-H4 (Cell Signaling, no. 2592, 1:1000), anti-NSD1 (Cell Signaling, no. 51076, 1:1000), Anti-V5-Tag (Cell Signaling, no. 13202S, 1:1000);

### Drug treatment

HAP1 cells were seeded in a 6-well plate (Corning, no. 3516) at 400,000 cells per well. After 24 hours, cells were washed with 1X DPBS (Gibco, no. 14190144) and treated with the indicated concentrations of dTAG-47 (Bio-Techne, no. 7530). Cells were harvested with 0.25% trypsin (Gibco, no. 15050065), washed with 1X DPBS then snap-frozen until further use.

### Co-immunoprecipitation (co-IP)

Endogenous co-IP was conducted with HAP1^dTAG-KDM5A^ whole cell extracts prepared with Pierce IP lysis buffer (Thermo Scientific, no. 87787) and supplemented with 5X Halt Protease and Phosphatase inhibitors (Thermo Scientific, no. 78441). Preclearing of the whole cell extracts with Pierce protein A/G beads (Thermo Scientific, no. 88802) was performed at 4 °C for 2 hours. Precleared extracts were then incubated with Pierce anti-HA magnetic beads (Thermo Scientific, no. 88836) overnight. Anti-HA magnetic beads were then washed with two times with cold IP wash buffer (50 mM HEPES pH 7.4, 150 mM NaCl, 5% glycerol and 0.2% NP-40) and eluted with 2 mg/mL HA peptides (GenScript, no. RP11735). To prepare western blot sample from co-IP eluates, 4X Laemlli sample buffer (Bio-Rad, no. 1610747) were added, and samples were boiled at 95 °C for 5 minutes.

### Statistical analyses

All data, if applicable, were presented as mean ± SD. Significant differences were determined by Student’s t-test. p < 0.05 was considered statistically significant.

## Acknowledgements

This research was supported by a grant from the Emerson Collective and the Helen Diller Family Comprehensive Cancer Center. Additional funding support was received from the UCSF Benioff Initiative for Prostate Cancer Research, the Breast Cancer Research Foundation, The Susan G. Komen Breast Cancer Foundation, NIH U54 CA209891, the V foundation for Cancer Research and the Gray Foundation. Figs. 5B was created using BioRender.com.

## Authors’ Contributions

H. Chen: Conceptualization, data curation, formal analysis, validation, investigation, visualization, methodology, writing–original draft, writing–review and editing. L. Sarah: Conceptualization, data curation, formal analysis, validation, investigation, visualization, methodology, writing–original draft, writing–review and editing. D. Pucciarelli: Data curation, validation, investigation. Y. Mao: Data curation, validation, investigation. M.E. Diolaiti: Formal analysis, supervision, methodology, writing–original draft, project administration, writing–review and editing. D.G. Fujimori: Conceptualization, resources, supervision, funding acquisition, writing–original draft, project administration, writing– review and editing. A. Ashworth: Conceptualization, resources, supervision, funding acquisition, writing–original draft, project administration, writing–review and editing.

## Supplementary Information Captions

**Supplementary Fig.1 Verification of HAP1^Δ5A^ and HAP1^Δ5Β^ cells.**

**A** Sanger sequencing traces showing frameshift mutations in HAP1^Δ5A^ and HAP1^Δ5Β^ cells.

**Supplementary Fig.2 Global analysis of ATAC-Seq data in HAP1^Δ5A^ and HAP1^Δ5Β^ cells.**

**A** Heatmaps showing the ATAC-seq merged peak regions in HAP1 (HAP-1 WT), HAP1^Δ5A^ (HAP-1 KDM5A-KO) and HAP1^Δ5Β^ (HAP-1 KDM5B-KO) cells.

**B** Heatmaps showing the distribution of ATAC-seq peaks at gene promoters (TSS) in HAP1 (HAP-1 WT), HAP1^Δ5A^ (HAP-1 KDM5A-KO) and HAP1^Δ5Β^ (HAP-1 KDM5B-KO) cells.

**C** Heatmaps showing the distribution of ATACseq peaks across gene bodies in HAP1 (HAP-1 WT), HAP1^Δ5A^ (HAP-1 KDM5A-KO) and HAP1^Δ5Β^ (HAP-1 KDM5B-KO) cells.

**Supplementary Fig.3 KRAB-ZNFs decreased H3K4me3 peaks in HAP1^Δ5A^ and HAP1^Δ5Β^ cells assayed by H3K4me3 ChIP-Seq.**

**A** Principal Component Analysis (PCA) showing variance in H3K4me3 distribution in HAP1 (HAP-1 WT) cells, HAP1^Δ5A^ (HAP-1 KDM5A-KO), HAP1^Δ5Β^ (HAP-1 KDM5B-KO) and CPI-455 treated HAP1 (HAP-1 CPI-455) cells. Duplicate samples for each condition are shown.

**B** Heatmaps showing the distribution of H3K4me3 peaks in HAP1 (HAP-1 WT), HAP1^Δ5A^ (HAP-1 KDM5A-KO), HAP1^Δ5Β^ (HAP-1 KDM5B-KO) and CPI-455 treated HAP1 (HAP-1 CPI-455) cells.

**C** Heat maps showing the distribution of H3K4me3 peaks at gene promoters (TSS) in HAP1 (HAP-1 WT), HAP1^Δ5A^ (HAP-1 KDM5A-KO), HAP1^Δ5Β^ (HAP-1 KDM5B-KO) and CPI-455 treated HAP1 (HAP-1 CPI-455) cells.

**Supplementary Fig.4 Enhanced ERVs transcription and immune response in KDM5A/B knockout cells.**

**A** Bar plots showing relative mRNA levels of the indicated ERV and ISG genes in HAP1 cells and CPI-455 treated HAP1 cells. Data are plotted as mean ± SEM.

**B** Bar plots showing the relative levels of the indicated ERVs and ISGs genes in K562 cells and CPI-455 treated K562 cells. Data are plotted as mean ± SEM.

**C** Histograms showing the relative levels of dsDNA in K562, K562^Δ5A^, K562^Δ5Β^ cells as assessed by FACS.

**D** Bar plots showing the relative mRNA levels of IFN-b and CXCL10 transcripts in HAP1 cells, treated with dsRNA, dsDNA or/and CPI-455. Data are plotted as mean ± SEM.

**Supplementary Fig.5 Uncropped blots for Fig. 1A**

The red rectangles outline the images used in the listed Figures

**Supplementary Fig.6 Uncropped blots for Fig. 3D**

The red rectangles outline the images used in the listed Figures

**Supplementary Fig.7 Uncropped blots for Fig. 4B**

The red rectangles outline the images used in the listed Figures

**Supplementary Fig.8 Uncropped blots for Fig. 4C**

The red rectangles outline the images used in the listed Figures

**Supplementary Fig.9 Uncropped blots for Fig. 5A**

The red rectangles outline the images used in the listed Figures

**S1 Table. Primers used for Real-time quantitative PCR.**

