## Supplementary tables and figures for "Histone demethylase enzymes KDM5A and KDM5B modulate immune response by suppressing transcription of endogenous retroviral elements"

**This PDF file includes:**

Supplementary Fig. 1 to 9
Supplementary Table 1

**Supplementary Fig.1 Verification of HAP1$^{\Delta5A}$ and HAP1$^{\Delta5B}$ cells.**

**A** Sanger sequencing traces showing frameshift mutations in HAP1$^{\Delta5A}$ and HAP1$^{\Delta5B}$ cells.

**Supplementary Fig.2 Global analysis of ATAC-Seq data in HAP1$^{\Delta5A}$ and HAP1$^{\Delta5B}$ cells.**

**A** Heatmaps showing the ATAC-seq merged peak regions in HAP1 (HAP-1 WT), HAP1$^{\Delta5A}$ (HAP-1 KDM5A-KO) and HAP1$^{\Delta5B}$ (HAP-1 KDM5B-KO) cells.

**B** Heatmaps showing the distribution of ATAC-seq peaks at gene promoters (TSS) in HAP1 (HAP-1 WT), HAP1$^{\Delta5A}$ (HAP-1 KDM5A-KO) and HAP1$^{\Delta5B}$ (HAP-1 KDM5B-KO) cells.

**C** Heatmaps showing the distribution of ATACseq peaks across gene bodies in HAP1 (HAP-1 WT), HAP1$^{\Delta5A}$ (HAP-1 KDM5A-KO) and HAP1$^{\Delta5B}$ (HAP-1 KDM5B-KO) cells.

**Supplementary Fig.3 KRAB-ZNFs decreased H3K4me3 peaks in HAP1$^{\Delta5A}$ and HAP1$^{\Delta5B}$ cells assayed by H3K4me3 ChIP-Seq.**

**A** Principal Component Analysis (PCA) showing variance in H3K4me3 distribution in HAP1 (HAP-1 WT) cells, HAP1$^{\Delta5A}$ (HAP-1 KDM5A-KO), HAP1$^{\Delta5B}$ (HAP-1 KDM5B-KO) and CPI-455 treated HAP1 (HAP-1 CPI-455) cells. Duplicate samples for each condition are shown.

**B** Heatmaps showing the distribution of H3K4me3 peaks in HAP1 (HAP-1 WT), HAP1$^{\Delta5A}$ (HAP-1 KDM5A-KO), HAP1$^{\Delta5B}$ (HAP-1 KDM5B-KO) and CPI-455 treated HAP1 (HAP-1 CPI-455) cells**.**

**C** Heat maps showing the distribution of H3K4me3 peaks at gene promoters (TSS) in HAP1 (HAP-1 WT), HAP1$^{\Delta5A}$ (HAP-1 KDM5A-KO), HAP1$^{\Delta5B}$ (HAP-1 KDM5B-KO) and CPI-455 treated HAP1 (HAP-1 CPI-455) cells**.**

**Supplementary Fig.4 Enhanced ERVs transcription and immune response in KDM5A/B knockout cells.**

**A** Bar plots showing relative mRNA levels of the indicated ERV and ISG genes in HAP1 cells and CPI-455 treated HAP1 cells. Data are plotted as mean ± SEM.

**B** Bar plots showing the relative levels of the indicated ERVs and ISGs genes in K562 cells and CPI-455 treated K562 cells. Data are plotted as mean ± SEM.

**C** Histograms showing the relative levels of dsDNA in K562, K562$^{\Delta 5A}$, K562$^{\Delta 5B}$ cells as assessed by FACS.

**D** Bar plots showing the relative mRNA levels of IFN-b and CXCL10 transcripts in HAP1 cells, treated with dsRNA, dsDNA or/and CPI-455. Data are plotted as mean ± SEM.

**Supplementary Fig.5 Uncropped blots for Fig. 1A**

The red rectangles outline the images used in the listed Figures

**Supplementary Fig.6 Uncropped blots for Fig. 3D**

The red rectangles outline the images used in the listed Figures

**Supplementary Fig.7 Uncropped blots for Fig. 4B**

The red rectangles outline the images used in the listed Figures

**Supplementary Fig.8 Uncropped blots for Fig. 4C**

The red rectangles outline the images used in the listed Figures

**Supplementary Fig.9 Uncropped blots for Fig. 5A**

The red rectangles outline the images used in the listed Figures

**S1 Table. Primers used for Real-time quantitative PCR.**

| Gene | F Primer | R Primer |
|---|---|---|
| *ZNF208* | TGGAGGAAGGAAAAGAGTCCTG | TCTCATGTCCACATTTTTCATACC |
| *ZNF676* | TCTTCCTGGGTATTGCTGCC | CTTCTATGCCCTGCTCTGGC |
| *HERVE* | GGTGTCACTACTCAATACAC | GCAGCCTAGGTCTCTGG |
| *HERVF* | CCTCCAGTCACAACAACTC | TATTGAAGAAGGCGGCTGG |
| *ERVV-2* | TCTGTTCCGCTCCAGGTTTC | CTGGGGAATCCCTCCTCAGA |
| *ERV3-1* | CATGGGAAGCAAGGGAACTAATG | CCCAGCGAGCAATACAGAATTT |
| *ERVW-1* | TTCACTGCCCACACCCAT | CCCCATCAGACATACCAGTT |
| *MAVS* | AGGAGACAGATGGAGACACA | CAGAACTGGGCAGTACCC |
| *IFIT1* | CTGAATGCAGCTCACCTCTG | GGATGGAATTGCCTGCTAGA |
| *IFIT3* | CTGAACTGCTCAGCCCACA | TCAGCTTGCCCTAAGCACTC |
| *IFN-β* | ACGCCGCATTGACCATCTATG | CGGAGGTAACCTGTAAGTCTGT |
| *CXCL10* | GGCCATCAAGAATTTACTGAAAGCA | TCTGTGTGGTCCATCCTTGGAA |
| *β-Actin* | GAGCACAGAGCCTCGCCTTT | TCATCATCCATGGTGAGCTG |

A

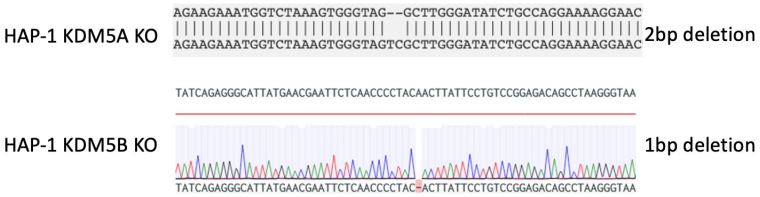


*Supplementary_Fig 1*

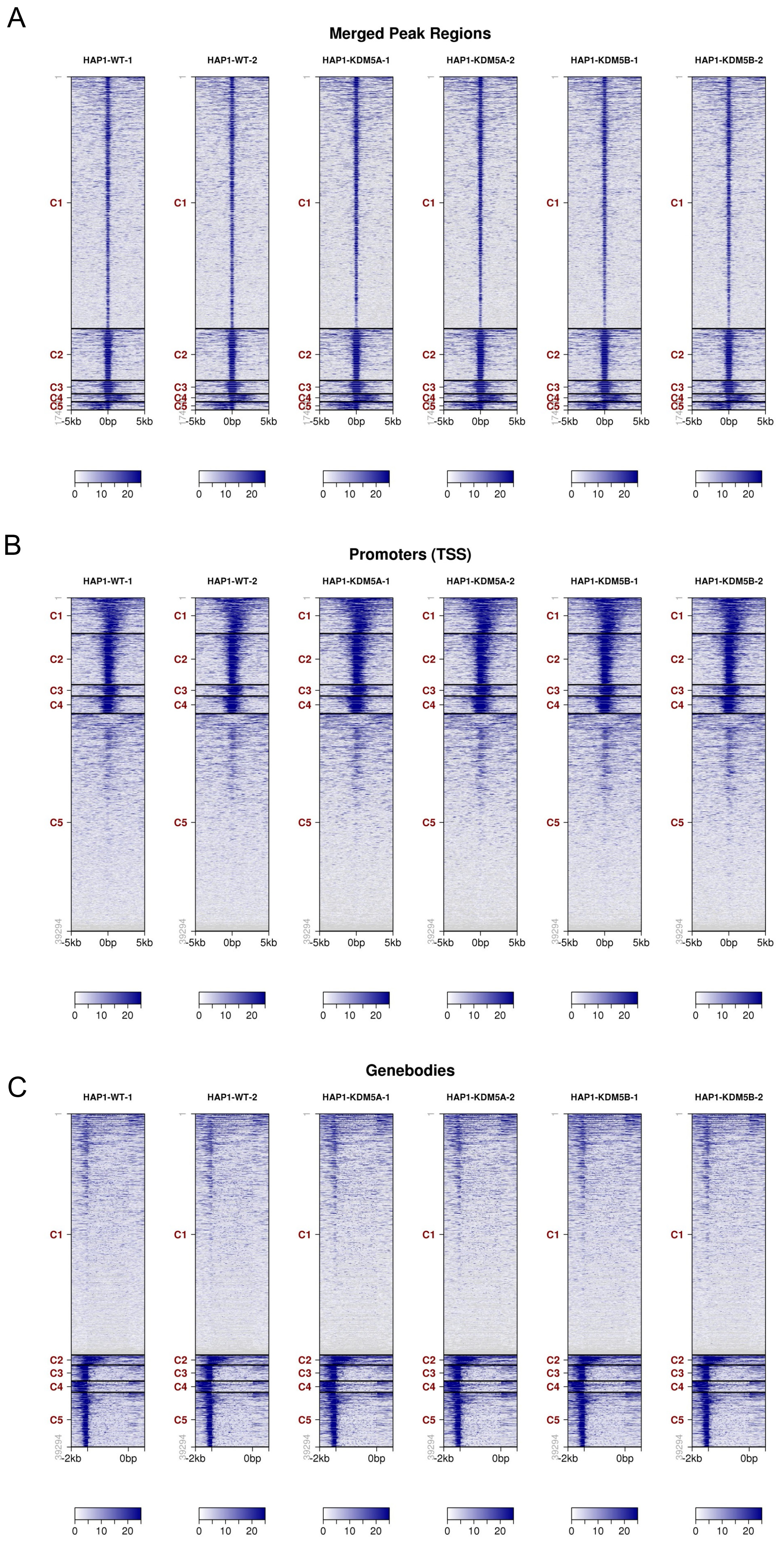


*Supplementary_Fig 2*

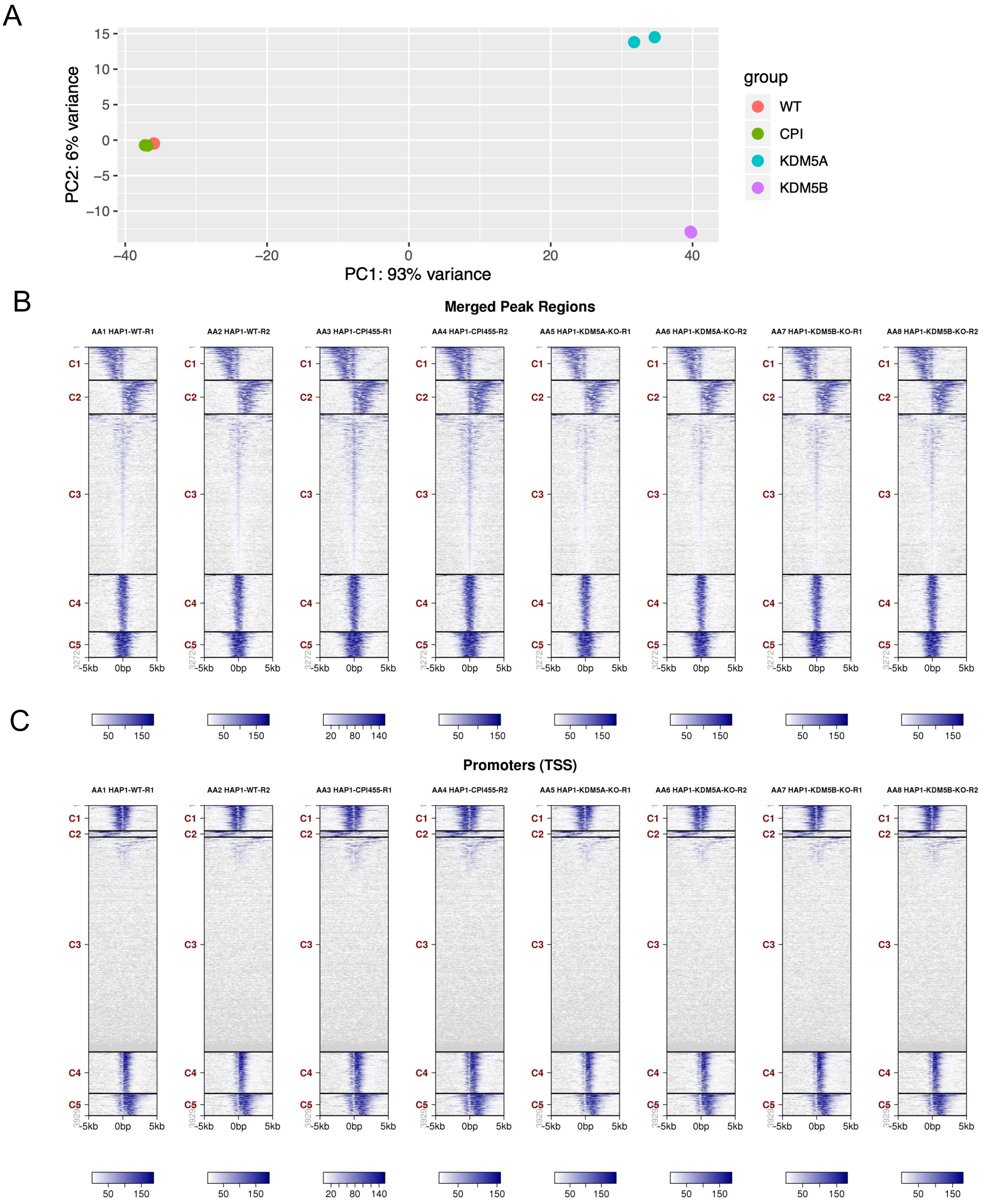


*Supplementary_Fig 3*

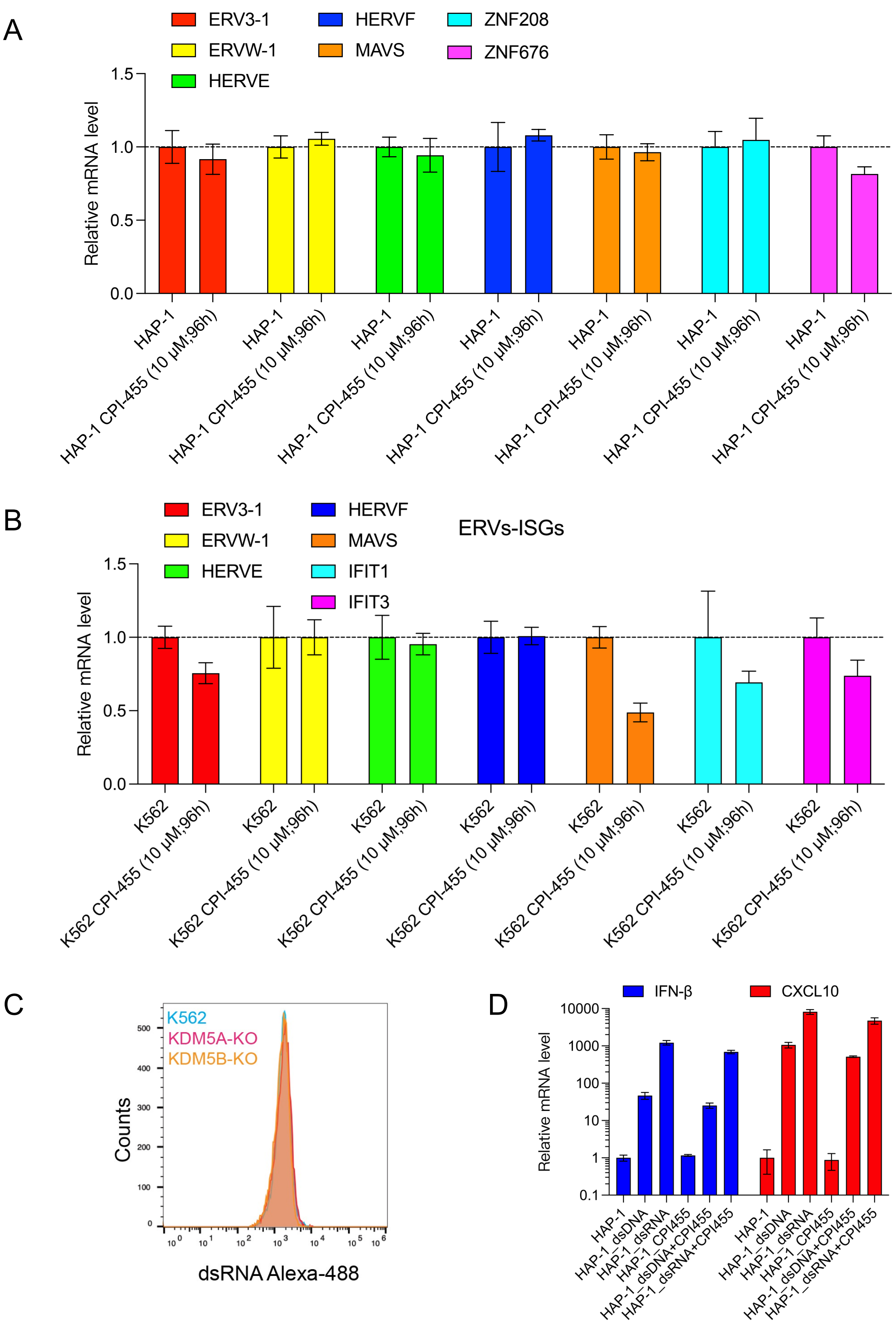


*Supplementary_Fig 4*

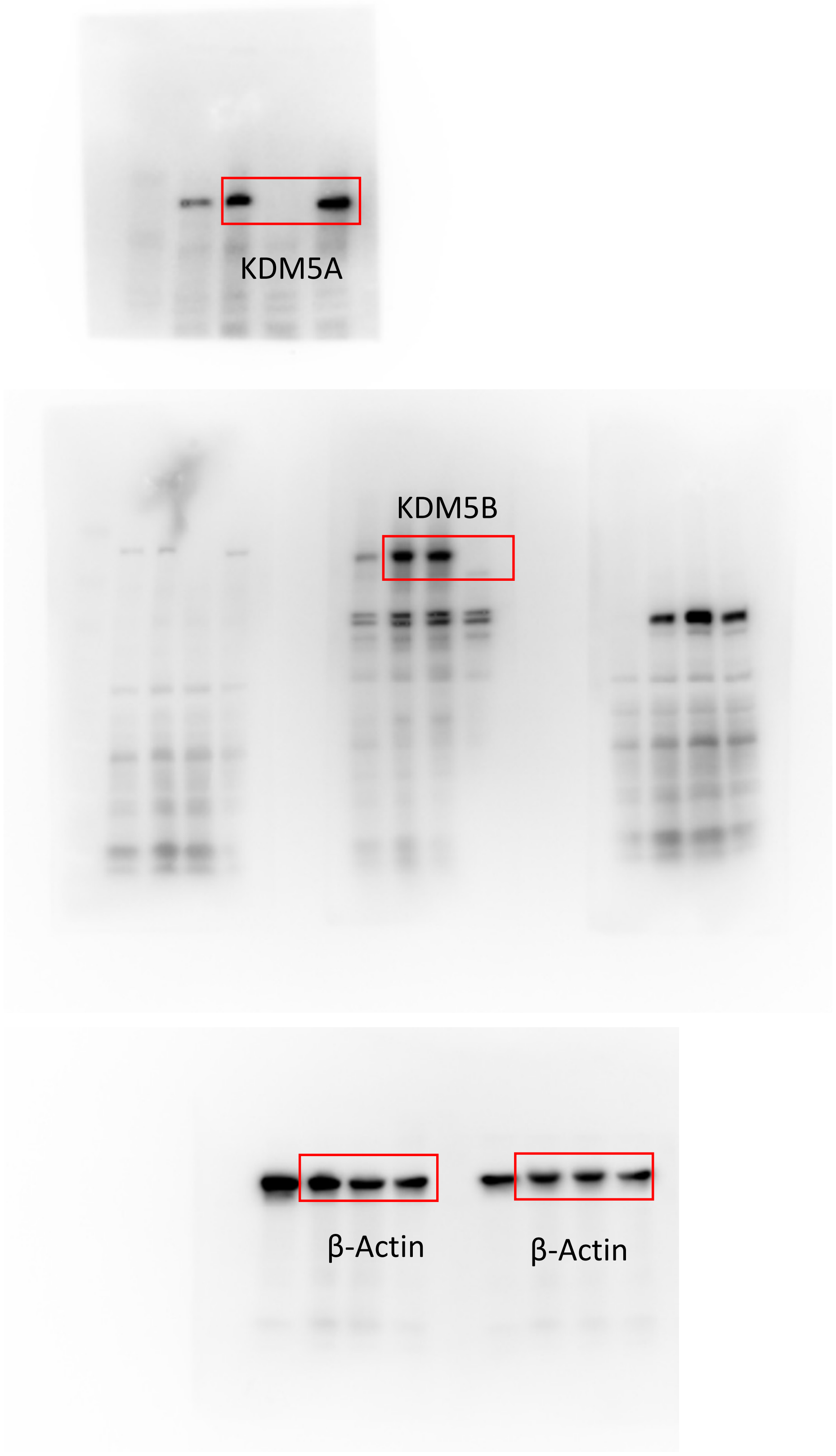


*Supplementary_Fig 5*

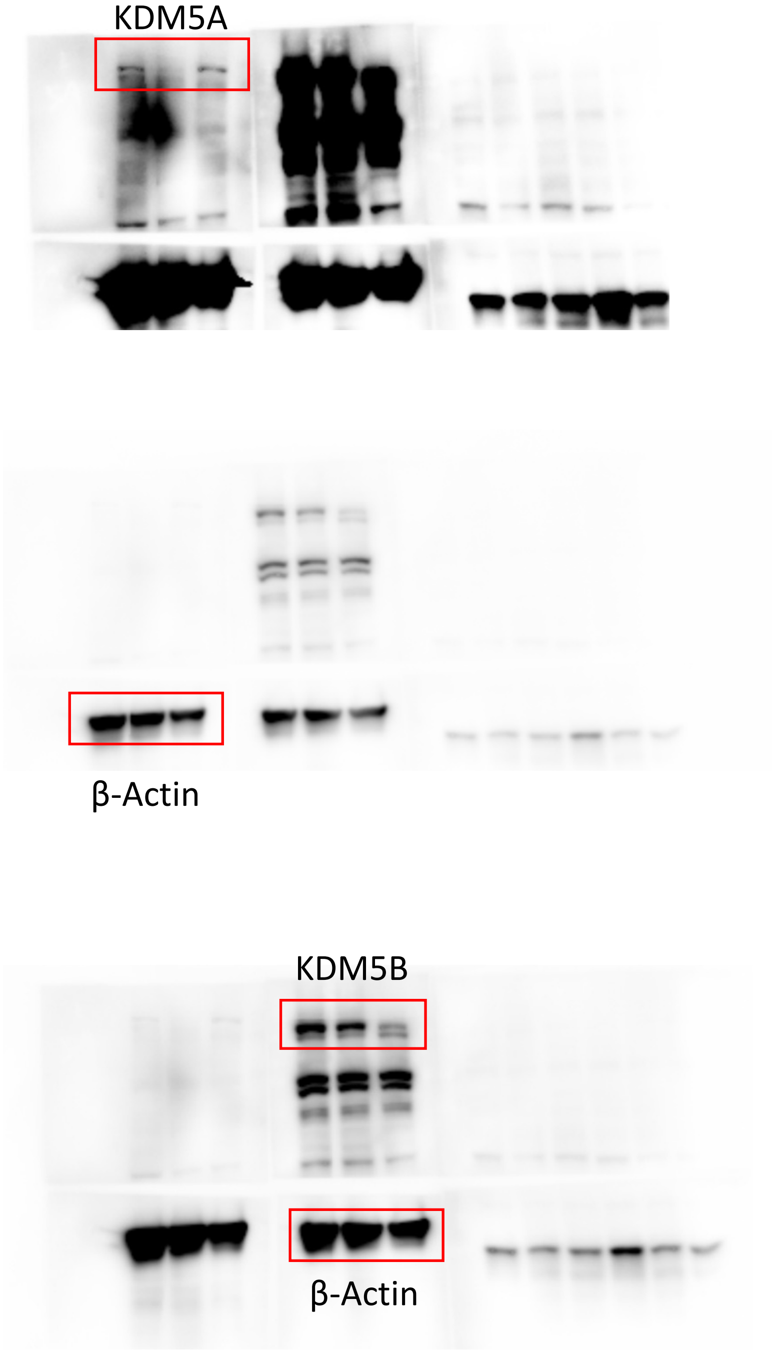


*Supplementary_Fig 6*

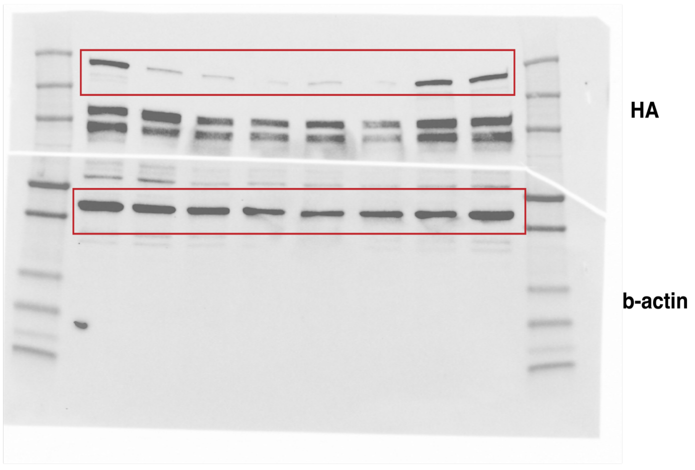

Western blots were done from the same SDS-PAGE gel

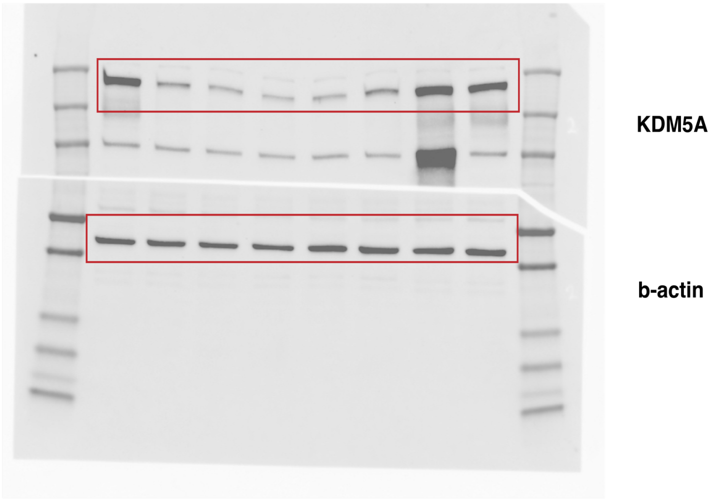

Western blots were done from the same SDS-PAGE gel

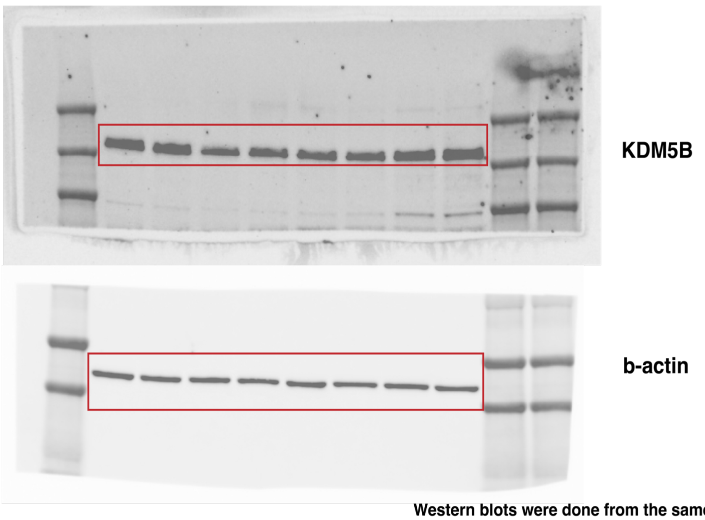

Western blots were done from the same SDS-PAGE gel

*Supplementary_Fig 7*

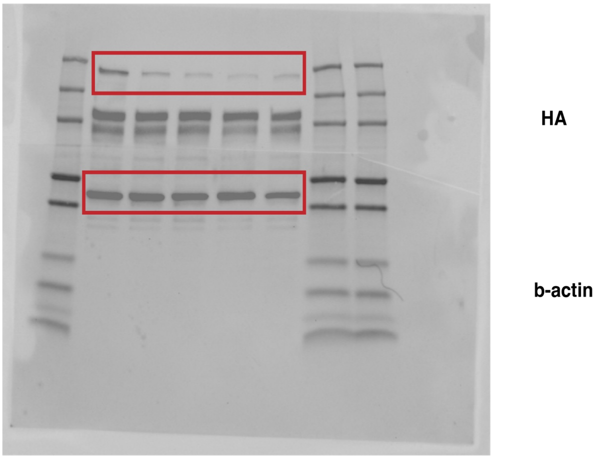


Western blots were done from the same SDS-PAGE gel

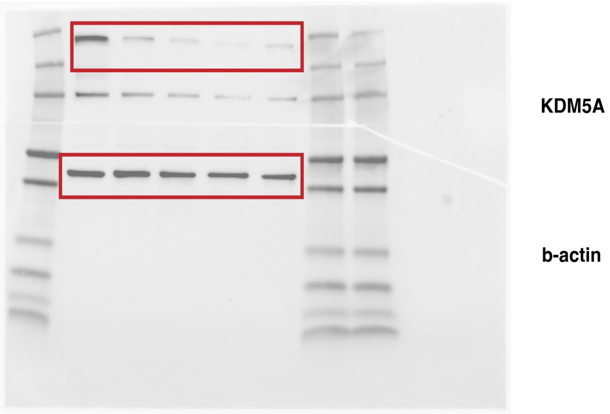


Western blots were done from the same SDS-PAGE gel

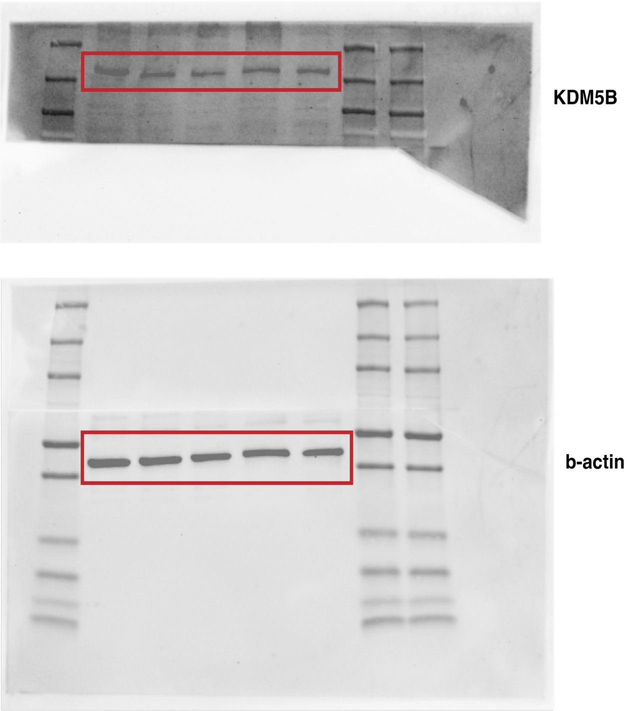


Western blots were done from the same SDS-PAGE gel

*Supplementary_Fig 8*

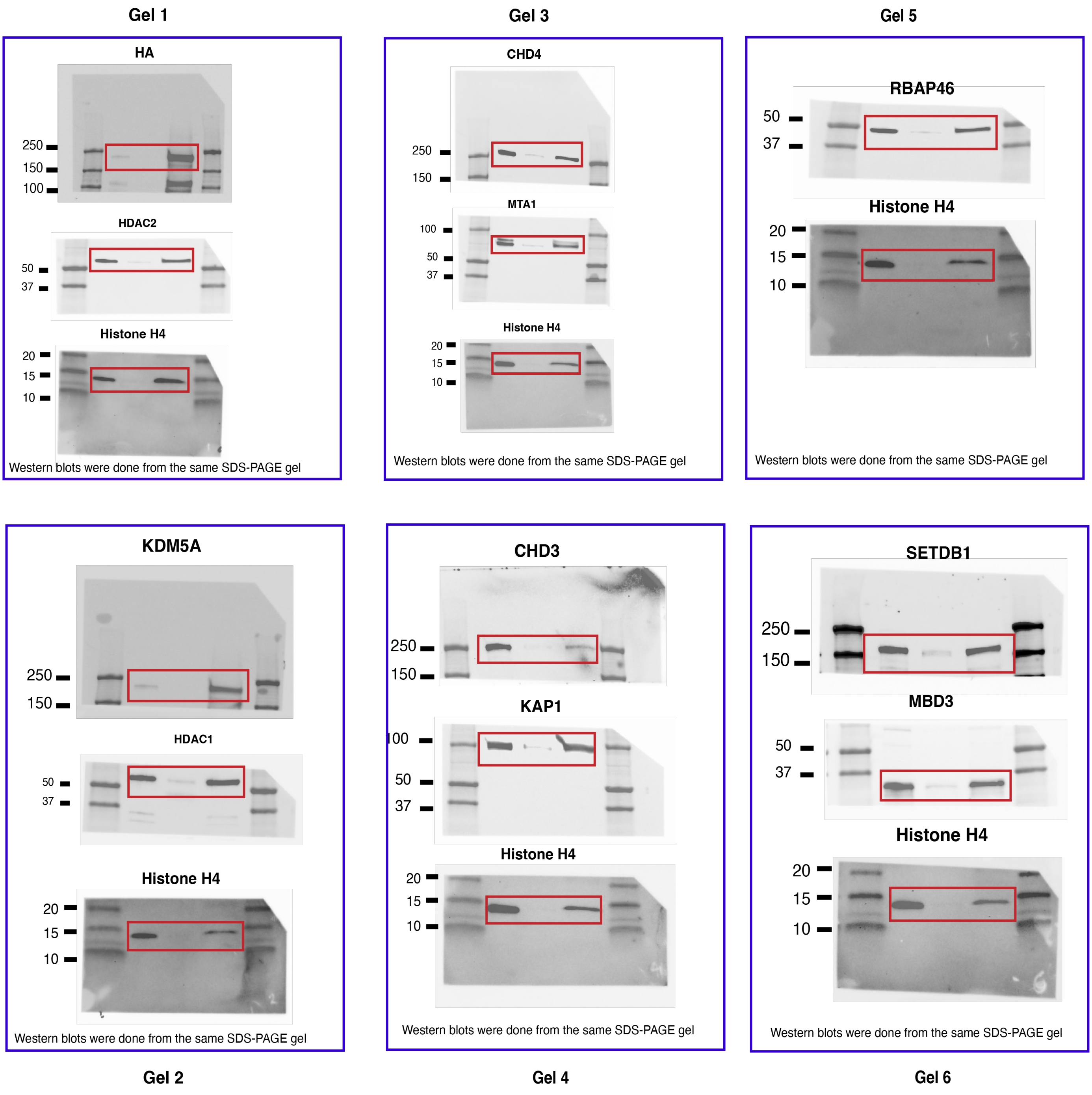


*Supplementary_Fig 9*
